# Structures of folding intermediates on BAM show diverse substrates fold by a uniform mechanism

**DOI:** 10.1101/2025.10.16.682720

**Authors:** Benjamin D. Thomson, Melissa D. Marquez, Shaun Rawson, Thiago M. A. dos Santos, Stephen C. Harrison, Daniel Kahne

## Abstract

The outer membranes of mitochondria, chloroplasts, and Gram-negative bacteria contain β-barrel membrane proteins that are assembled by conserved multi-subunit machines. In bacteria, the β-barrel assembly machine (BAM) folds over a hundred compositionally different substrates into barrels that vary greatly in size. Some larger barrels require globular proteins to plug the barrel lumen. How a single machine can assemble such different barrels is unknown. Here we report three structures representing progressively folded stages of a 16-stranded barrel engaged with BAM, as well as the structure of a late-stage folding intermediate of a 26-stranded substrate folding around its soluble lipoprotein plug on BAM. We find that BAM catalyzes folding of these substrates by a uniform mechanism in which BAM undergoes major distortions to accommodate the nascent barrel.

## Main

Membrane proteins with a β-barrel topology are present only in the outer membranes of mitochondria, chloroplasts, and Gram-negative bacteria (*1*, *2*). They are folded by a conserved protein machine (*3–7*). In *Escherichia coli*, the β-barrel assembly machine (BAM) consists of a β-barrel protein, BamA, and four lipoproteins (*4*, *8–14*) (Fig. 1A). Of the five subunits, only BamA (BamA^M^) is required to catalyze folding (*8*, *15*, *16*). Structures of substrate-engaged BAM complexes show that the C-terminal end of BAM substrates form a hydrogen-bonded β-strand to the N-terminal end of BamA^M^ (*17–23*). These structures suggest that barrels fold in a C-to-N direction by addition of β-strands to a template strand, with BamA^M^ providing the first template strand (*17*, *18*, *24*).

**Fig. 1:**
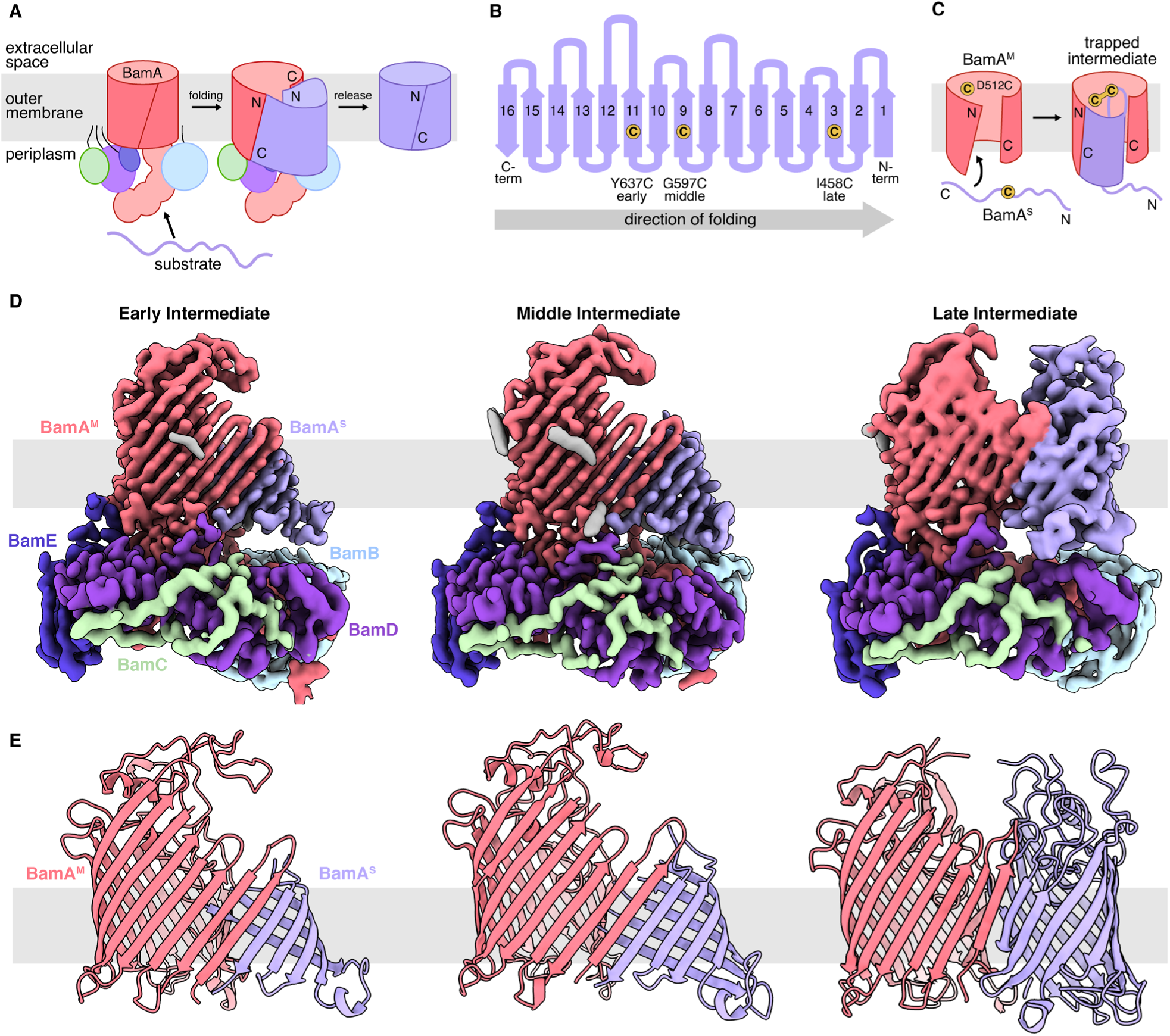
Trapping BAM in native intermediate states results in progressively folded substrate. (**A**) Summary of BAM-mediated β-barrel assembly. Substrates are held by the N-terminal strand of BamA^M^ in a hybrid barrel sheet before release from the machine. (**B**) Topological depiction of the BamA^S^ substrate barrel. Arrow shows direction of folding of the BamA^S^ β-barrel domain from C- to N-terminal strand. Positions of three engineered cysteines that crosslink to BamA^M^(D512C) in native folding intermediates are shown as yellow circles. (**C**) Scheme of substrate disulfide crosslinking to the BamA^M^ luminal wall. Cysteine-containing substrate strands with long residence times within the BamA^M^ lumen make a disulfide crosslink with BamA^M^(D512C), forming a trapped intermediate complex. (**D**) Cryo-EM structures of Early, Middle and Late Intermediates of BamA^S^ folding on BAM, left to right. Displayed maps were postprocessed using DeepEMhancer. A composite map is shown for the Late Intermediate. Densities corresponding to BamA^M^, BamB, BamC, BamD, BamE, and BamA^S^ are colored in salmon, light blue, light green, purple, dark blue, and lavender respectively. From left to right, more and more folded substrate density is present. (**E**) BamA^M^ and BamA^S^ barrel regions of models built into cryo-EM structures of the Early, Middle and Late Intermediate complexes.

BAM folds an array of different substrates that vary greatly in size (from 8 to 26 β-strands in circumference) and shape (*25–27*). The most well-studied substrate is EspP, a small 12-stranded barrel, and the majority of studies on BAM folding have focused on this substrate (*18–20, 22*). As barrels get larger, their assembly must become more complex. In fact, many large barrels contain soluble plugs which seal the barrel lumen (*26–32*). For example, LptD, the barrel component of the outer membrane translocon that moves lipopolysaccharides into the outer membrane, contains the lipoprotein LptE (*26*, *27*, *33*); LptD does not assemble without LptE (*34–36*). How a single machine can fold a wide array of structurally diverse substrates is the question we have sought to answer here.

We recently detected stable folding intermediates of two native barrel substrates, BamA itself (BamA^S^) and LptD, during folding on BAM *in vivo* (*37*). We have now isolated three folding intermediates of BamA^S^ engaged with BAM and obtained a series of structures that show how folding of an individual substrate barrel proceeds. We have also determined the structure of a late-stage folding intermediate of the 26-stranded LptD folding around its LptE plug while attached to BAM. These new structures show that BAM catalyzes folding of many different substrate barrels by a uniform mechanism in which BamA^M^ undergoes major distortions to accommodate the nascent barrel.

### Progressively Folded Intermediates

We previously used disulfide crosslinking to detect stable intermediates during folding of a substrate barrel on BAM (*37*) (Fig. 1B and C, Extended Data Fig. 1). We substituted a cysteine (D512C; Fig. 1C) facing the lumen of machine BamA^M^. We expressed this BamA^M^ together with an array of different, folding-competent, full-length BamA^S^ substrate barrels, each with a cysteine placed in one of the 16 strands of the BamA^S^ barrel (Fig. 1B). The longer a particular substrate strand spends in the lumen of BamA^M^, the more likely it is to become trapped by forming a disulfide bond with the luminal cysteine (Fig. 1C). Although the residence times of most substrate strands were too short for crosslink formation, BamA^S^ with cysteine in β-strands 12-8 and 3 crosslinked strongly to the cysteine in the machine lumen. We concluded that BamA^S^ folding is slow enough at these specific intermediate states in substrate assembly for disulfide crosslinks to form efficiently *in vivo*, offering the opportunity to determine structures for distinct stages of folding of a single substrate.

If substrates fold in a C-to-N-terminal direction on BAM, the structure of BAM-BamA^S^ (G597C) should represent a later folding intermediate than that of BAM-BamA^S^ (Y637C) and BAM-BamA^S^ (I458C), should represent a yet later one. We overexpressed and purified these Early, Middle, and Late Intermediate complexes and determined their structures by cryogenic electron microscopy (cryo-EM). We obtained structures at final reconstructions of 3.6Å, 3.3Å, and 3.9Å, respectively (Fig. 1D, Extended Data Fig. 2,3,4, Extended Data Table 1). Each map had density corresponding to BamA^M^BCDE, and BamA^S^, with progressively more BamA^S^ substrate density in the sequence from the Early complex to the Middle and Late ones (Fig. 1D lavender density).

We built atomic models into each density using previously published structures (*10*, *11*, *17*) (Fig. 1E, Extended Data Fig. 5). In each intermediate, BamA^M^ and BamA^S^ form an asymmetric hybrid barrel, with the N-terminal edge of the BamA^S^ sheet facing into the extended, aqueous lumen. Substrate β-strands 16-12, 16-10, and 16-4 are present in the Early, Middle Late Intermediate structures, respectively, showing that when folding on BamA^M^, β-strands of BamA^S^ add progressively to the emerging substrate barrel.

### Path of Substrate Strands

To control for the possibility that the luminal disulfide crosslink we used to trap these folding intermediates biased the structures toward nonnative conformations, we determined whether features present in the structures could be detected *in vivo* without the luminal crosslink.

In the structure of the Early Intermediate, BamA^S^ β-strands 16-12 are present as a sheet that curves from the hybrid barrel interface and forms a back-to-back lateral seal against BamA^M^ β-strand 16 (Fig. 2A and B). To test whether this back-to-back interaction occurs in the absence of a luminal crosslink (D512C), we carried out a separate *in vivo* disulfide crosslinking experiment, by substituting a single cysteine at position A710, M711 or A712 in β-strand 12 of BamA^S^. In the Early Intermediate structure, the periodic beta sheet allows A710 and A712 to face the BamA^M^ C-terminal strand and M711 to face the lumen (Fig. 2C). We expressed each of these three BamA^S^ substrate variants together with a BamA^M^ variant containing I806C in BamA^M^ β-strand 16 and then assessed BamA^M^- BamA^S^ crosslink formation by pulldown and immunoblotting. We found that alternating residues in BamA^S^ strand 12 crosslinked to BamA^M^ I806C; A710C and A712C formed much stronger crosslinks than did M711C (Fig. 2D). Thus, we can detect the features observed in the Early Intermediate structure in the absence of a luminal disulfide crosslink at BamA^M^ (D512C).

**Fig. 2:**
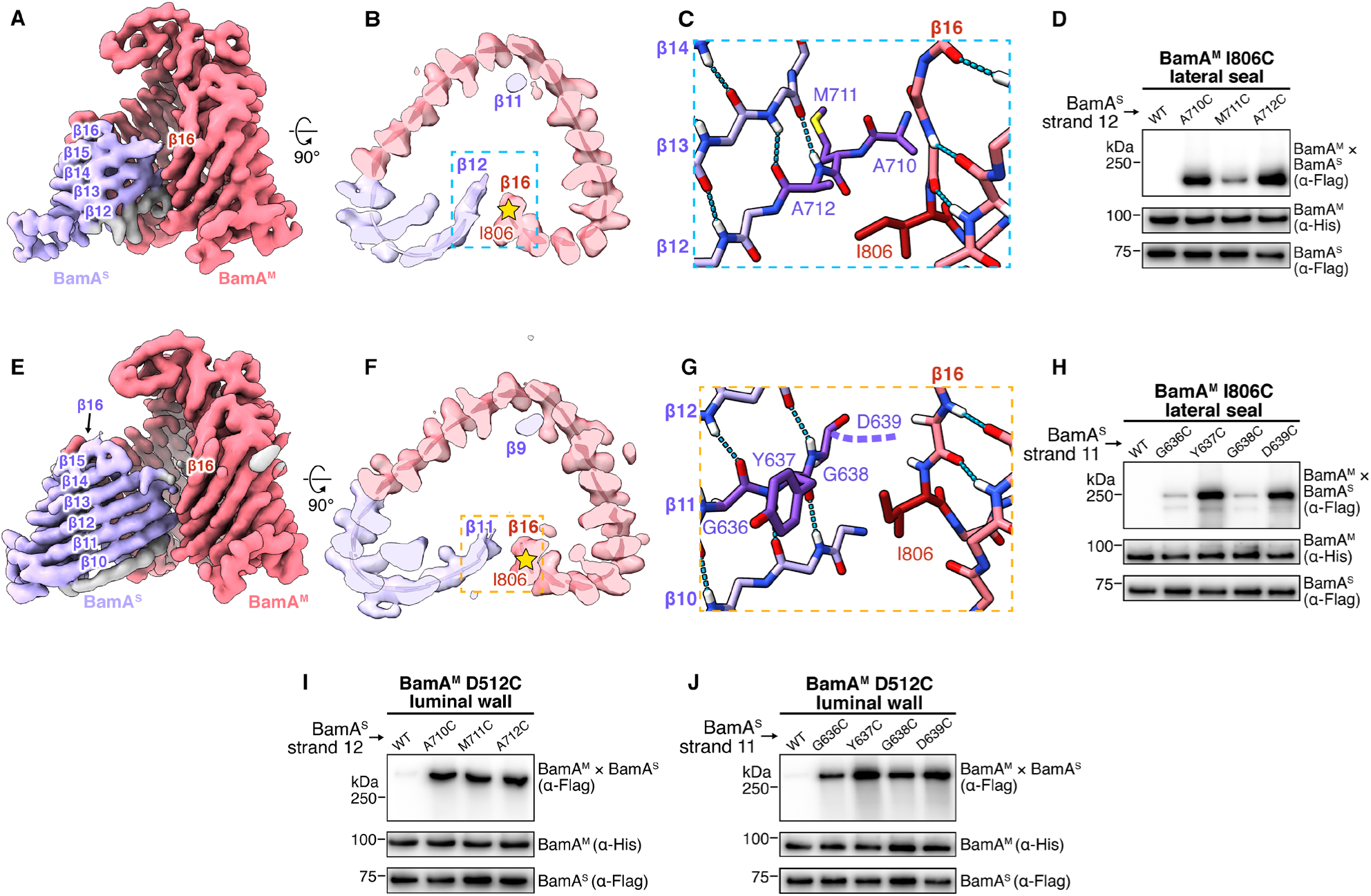
β-strands are disordered in the BamA^M^ lumen and fold *in vivo* at the edge of the substrate. (**A**) Early Intermediate barrel density turned 180° from the view in Fig. 1D. BamA^S^ β-strands 16-12 are labelled. (**B**) In-membrane cross-section of Early Intermediate hybrid barrel. β-strand 11 is trapped in the BamA^M^ lumen. I806 in BamA^M^ is labeled (yellow star). The back-to-back lateral seal between BamA^M^ and BamA^S^ is indicated by blue, dashed-line box. (**C**) Early Intermediate back-to-back interaction. Side chains of BamA^S^ A710 and A712 face I806 of BamA^M^, while M711 faces the lumen. (**D**) Disulfide crosslinking of BamA^S^ β-strand 12 to BamA^M^ I806C assessed by α-FLAG immunoblotting of His-purified samples (upper row). α-His immunoblots provide a pulldown efficiency control (middle). α-FLAG immunoblots of cell lysates provide a BamA^S^ expression control (lower). (**E**) Middle Intermediate barrel density. BamA^S^ β-strands 16-10 are folded. (**F**) Cross-section of Middle Intermediate hybrid barrel. The back-to-back lateral seal is indicated by the orange, dashed-line box. (**G**) Middle Intermediate back-to-back interaction. BamA^S^ Y637 faces I806 of BamA^M^. BamA^S^ D639 is not resolved (dashed line) but faces outward in folded BamA^S^. G636 and G638 face the lumen. (**H**) Disulfide crosslinking of BamA^S^ β-strand 11 to BamA^M^ I806C. (**I, J**) Disulfide crosslinking of tested residues to BamA^M^ lumen D512C.

In the structure of the Middle Intermediate, the β-strand that comes closest to BamA^M^ I806 is BamA^S^ β-strand 11 (Fig. 2E and F). Following the approach just described for the Early Intermediate, we substituted an individual cysteine at either position G636, Y637, G638, or D639 in β-strand 11 of BamA^S^ (Fig. 2G), and expressed each of these four BamA^S^ substrate variants together with BamA^M^ (I806C). Crosslinking of these residues to BamA^M^ I806C alternated, with Y637C and D639C crosslinking more strongly than G636C or G638C (Fig. 2H). Although BamA^S^ D639 is not resolved in the Middle Intermediate structure, it faces outward in the folded BamA barrel (Fig. 2G dashed purple line, Extended Data Fig. 6A and B). Therefore, like the Early Intermediate, the Middle Intermediate has features that can be detected in the absence of the luminal crosslink used initially to identify the intermediates *in vivo* (*37*).

We have reported many substrate interactions with different locations on the BamA^M^ luminal wall (*37–39*). Next to BamA^M^ D512C in the present structures, we see only relatively weak density, which we suggest corresponds to the crosslinking substrate cysteine (Fig. 2B and F, Extended Data Fig. 6C and D). The absence of strong features suggests that, while substrate strands do interact with the BamA^M^ lumen during folding, they do not do so as ordered β-strands. We assessed crosslinking of A710C, M711C or A712C in BamA^S^ strand 12, as well as G636, Y637, G638, or D639 in BamA^S^ strand 11, to the luminal wall BamA^M^(D512C). Within each strand, each residue crosslinked to the BamA^M^ luminal wall with approximately equal intensity (Fig. 2I and J), rather than in the alternating pattern expected of an ordered strand, indicating that substrate strands interact with the luminal wall in a less ordered conformation than the one in which they interact with the open edge of the BamA^M^ C-terminal strand at the lateral seal. Thus, we have detected two substrate strands (β-strands 12 and 11) in two distinct states – one in which the strand is disordered in the lumen, and the other in which the strand is ordered at the lateral seal. We infer that substrates enter the lumen as unfolded polypeptide and fold at the lumen-facing, N-terminal edge of the folded substrate chain.

The cryo-EM density for the Middle Intermediate contains extra density associated with a portion of the BamA^S^ β-strand 10 (Fig. 2E grey density, Extended Data Fig. 7), but the strand is not clearly interpretable as a β-strand of specific sequence. Because the next strand to be folded in the process is crosslinked to the BamA^M^ luminal wall, this density is likely not part of the on-pathway process. A similar feature is present in the Early Intermediate structure. These densities suggest that in the trapped complex, nascent substrate polypeptide within the BamA^M^ lumen can form partial or transient β-strands with the open edge of the folded barrel.

### Variable Contortion of BamA^M^

The observation that disulfide crosslinking can be used to observe unfolded substrate strands in the BamA^M^ lumen indicates that the disulfide crosslinking method could report on the lifetime of a specific stage of the folding process. If nascent substrate strands add sequentially from the lumen as our structures suggest, then intervening loops that connect adjacent strands will influence the rate and efficiency of folding from the lumen.

To test this consequence of a sequential folding mechanism, we deleted loop 5 (ΔL5) from BamA^S^ and assessed the crosslinking of proximal BamA^S^ strands 8, 9, and 10 to BamA^M^ (D512C) in the lumen of the complex (Fig. 3A and B). Crosslinking of each of the strands was greater for BamA^S^ ΔL5 than for wild-type BamA^S^. This result suggests that L5 influences the local structure of β-strands 9 and 10 during folding. As a control, we performed the same experiment with loop 1 (ΔL1), which lies far from strands 8-10 of the substrate. We found that ΔL1 had no effect on crosslink formation. These results suggest that changing local structure can change the time that strands spend in an unfolded state within the BamA^M^ lumen and that our disulfide crosslinking method can report on distinct stages of folding.

**Fig. 3:**
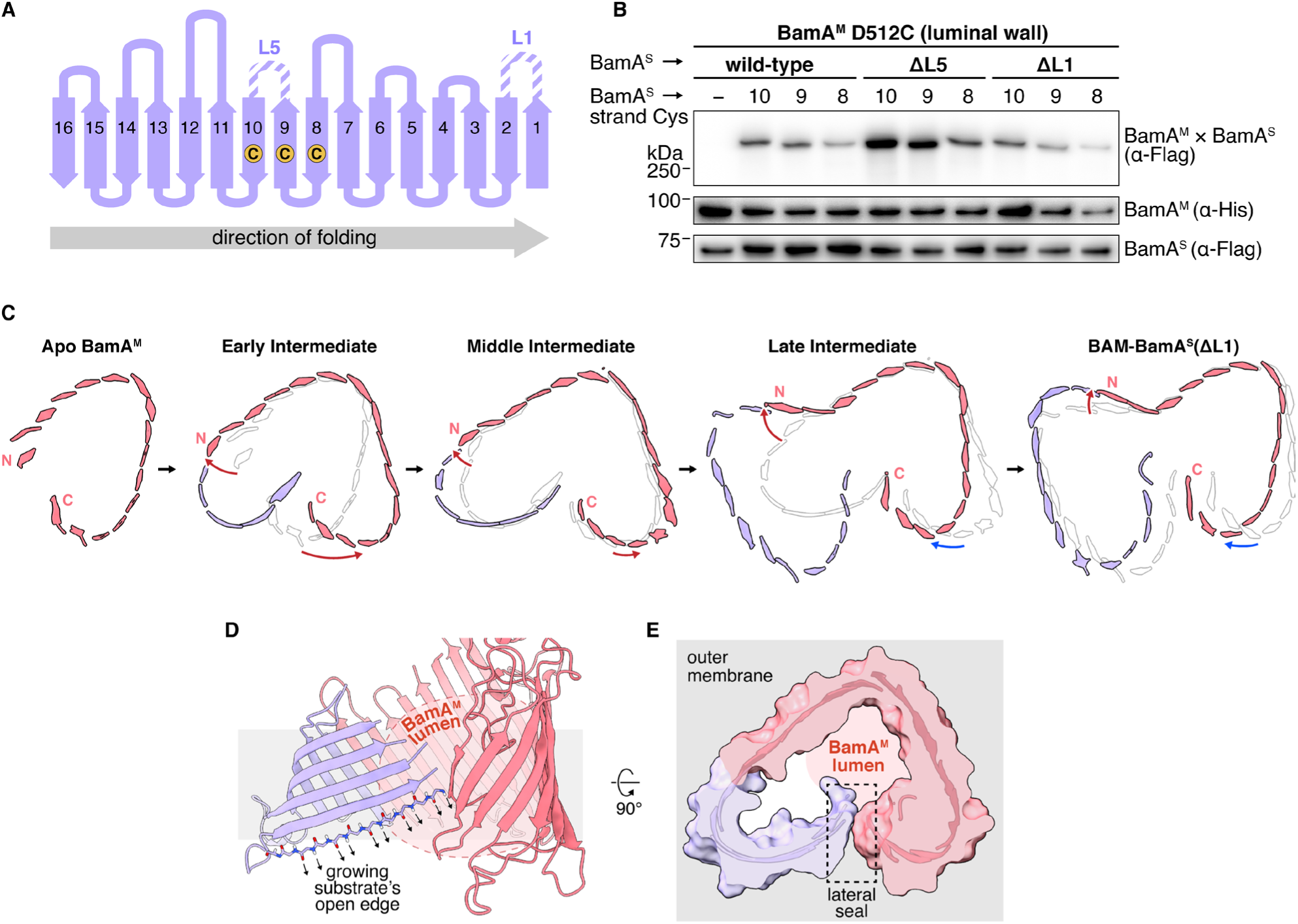
BamA^M^ variably distorts throughout folding of a single substrate. (**A**) Topological map of the BamA^S^ barrel showing positions of deleted extracellular loops 5 and 1 and locations of the cysteines in β-strands 8, 9, or 10 used for testing luminal disulfide crosslinking (yellow circles). (**B**) Disulfide crosslinking of BamA^S^ β-strands 8, 9, and 10 to the machine luminal wall in the presence and absence of loop deletions ΔL1 and ΔL5. Results are laid out as described in Fig. 2D. (**C**) In-membrane cross sections of BamA^S^ folding intermediate structures aligned at BamA^M^ β-strands 4-6. To illustrate displacements during folding, each intermediate is overlaid on the structure of the previous step (colored on light grey, PDB: 5LJO). Red arrows show flexion of C- and N- terminal parts of the curved sheet away from the native structure of BamA^M^, as superposed on strands 4-6. Blue arrows show relaxation back toward the native position. (**D**) Middle Intermediate barrel region enlarged to illustrate the direction in which the N- terminal backbone hydrogen bonding atoms project throughout the folding process. (**E**) In-membrane cross-section of a surface model of the Middle Intermediate structure demonstrating the back-to-back lateral seal that is present throughout folding. View is rotated 90° from (D). The BamA^M^ lumen and outer membrane are highlighted.

Fig. 3 shows alignments of the in-membrane cross-sections of our progressively folded structures as well as of our previously published BAM-BamA^S^(ΔL1) structure (*17*) and of apo BamA^M^ (*10*). Each structure is overlaid on that of the prior snapshot, displayed as a ghost (Fig. 3C). The alignments, superposed using BamA^M^ β-strands 4-6, show how the machine barrel distorts continuously depending on the specific stage of substrate folding.

The displacements of portions of the barrel N- and C-terminal strands relative to the superposition zone (strands 4-6) may help explain why stable folding intermediates can be detected at certain stages, but not at others (*37*). In the early stages of folding, the two displacements of both parts of the BamA^M^ barrel work together to open the barrel progressively wider (compare Fig. 3C, Apo BamA^M^, Early and Middle). The Middle Intermediate, in which the BamA^M^ C- and N-terminal strands are approximately 30Å apart, is the widest open state yet observed. The accompanying strain may account for the kinetics that have allowed us to observe an early set of long-lived folding intermediates *in vivo* during folding of BamA^S^ β-strands 12-8 (*37*) as well as distinct unfolded and folded states for BamA^S^ β-strands 11 and 12 (Fig. 2). As folding progresses through the Late Intermediate, the BamA^M^ N-terminal strands continue to flex, acquiring a severe outward bend, coupled to relaxation of the C-terminal region back toward its native state (Fig. 3C, compare Middle Intermediate, Late Intermediate, BAM-BamA^S^ (ΔL1)). Recent work has demonstrated that inward-facing glycines within membrane β-barrels favor tight inward curvature of the β-sheet (*40*). Progressive generation of the outward N-terminal strain, in a region rich in inward-facing glycines (Extended Data Fig. 8), might account for the kinetics that have allowed us to observe the Late Intermediate *in vivo*. The energetic benefit of progressive formation of substrate β-sheet probably pays the energetic cost of distorting BamA^M^.

Throughout the folding process, one edge of the substate participates in a hybrid sheet, with the other edge free and facing the BamA^M^ lumen, to facilitate addition of unfolded strands (Fig. 3D). At each stage in folding, there is a hydrophobic seal between the BamA^M^ C-terminal strand and the surface of the newly formed substrate sheet (Fig. 3E). The surface region of the BamA^S^ barrel that makes this seal changes as folding progresses, while maintaining seal integrity (Fig. 3C). In each stage of folding we have captured, BamA^M^ distorts from the state observed in apo BamA^M^ in directions such that relaxation of the outward flex would generate a force in just the direction needed to hold the nonspecific seal closed as folding progresses, while allowing the two opposing surfaces to slide across each other.

### The Late Intermediate Accumulates when Release Slows

Unlike the BamA^M^ barrel which distorts throughout folding to accommodate growth of the substrate barrel (Fig. 3C), BamA^S^ maintains the curvature of the native barrel as substrate strands fold (Fig. 4A and B, Extended Data Fig. 9). Release of a folded barrel from BAM requires exchange of hydrogen bonds between the C-terminal strand of the substrate and the N-terminal strand of BamA^M^. This exchange can occur only when the C- and N-terminal strands of the substrate barrel come sufficiently close together (*17*, *20*). Deleting loop 1 from BamA^S^ impairs release (*17*). If altering the substrate structure far from the N-terminal strand also impairs release, the intensity of the disulfide crosslink formed between the Late Intermediate and BamA^M^ will increase. We tested this prediction, first verifying that deleting either loop 1 (ΔL1) or periplasmic turn 1 (ΔT1) from BamA^S^ (I458C) (Fig. 4C, purple) increased the disulfide crosslinking intensity (Fig. 4D), to show that we can detect slow-releasing substrates by monitoring buildup at the Late Intermediate folding stage. We then deleted loops 3 (ΔL3) or 4 (ΔL4) from BamA^S^ (I458C) and assessed crosslinking to BamA^M^ (D512C). We chose these loops because by the Late Intermediate stage, they have the same conformation as in native BamA (Fig. 4B and C, blue). Like ΔL1 and ΔT1, both loop deletions enhanced the crosslinking intensity (Fig. 4E). Thus, these loop deletions do not prevent BamA^S^ from forming a Late Intermediate, but slow release from BAM.

**Fig 4:**
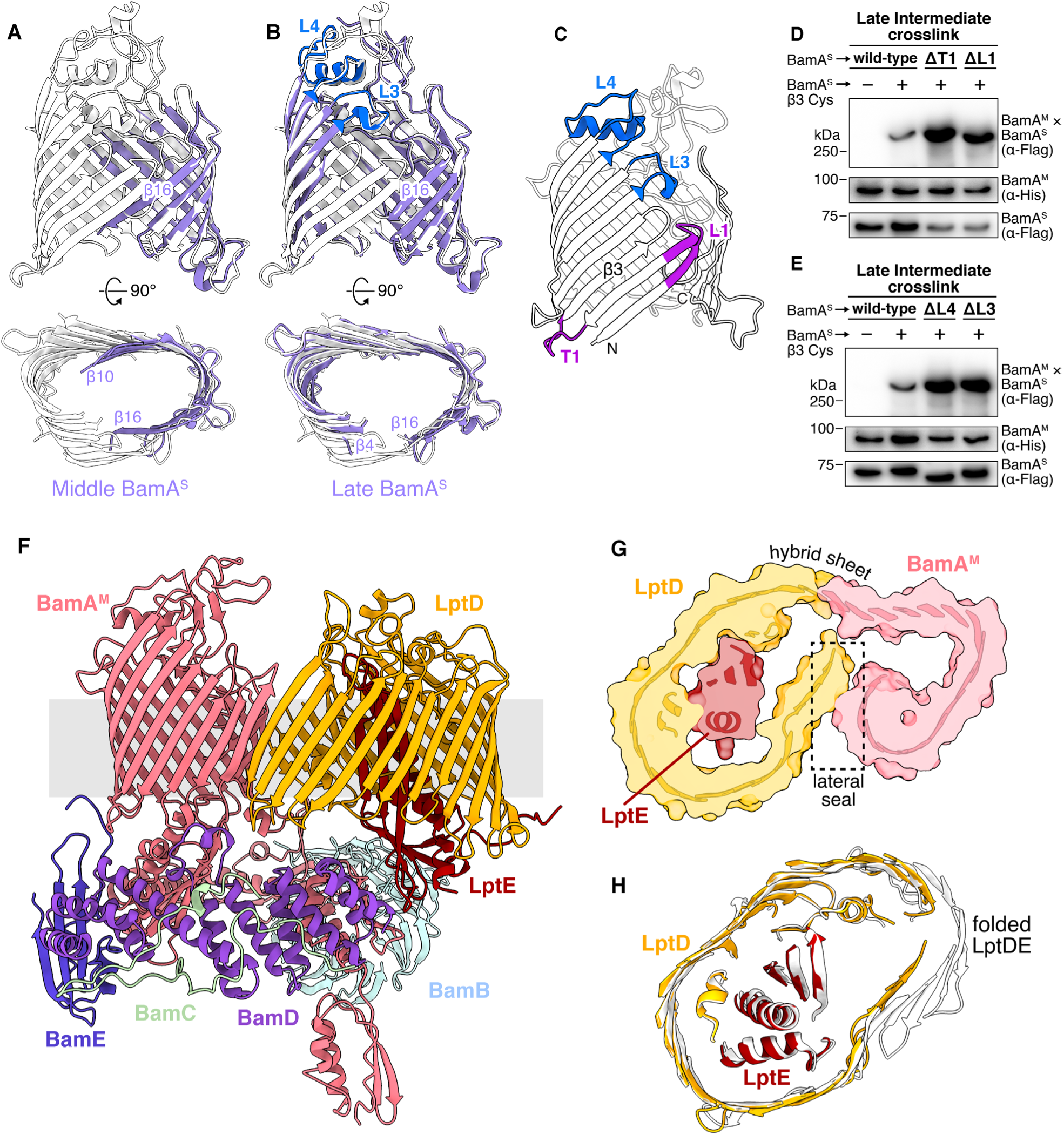
Features of BamA^S^ folding are conserved in LptD folding. (**A, B**) Overlay of the closed folded BamA barrel (white, PDB: 5D0O) with the BamA^S^ substrate (lavender) in the Middle Intermediate (A) and Late Intermediate (B). Head-on views (upper) and in-membrane cross-sections (lower) are shown. (**C**) Positions of periplasmic turn 1 (T1), and extracellular loops 1 (L1), 3 (L3), and 4 (L4) within the folded BamA barrel. L1 and T1 (purple) are proximal to the N-terminal exchanging strand. L3 and L4 (blue), are distant from the seam. (**D**) Disulfide crosslinking in the Late Intermediate (BamA^S^ I458C in β-strand 3 to BamA^M^ D512C) with or without deletions ΔT1 and ΔL1. Data are laid out as described in Fig. 2D. (**E**) Disulfide crosslinking in the Late Intermediate with or without deletions ΔL4 and ΔL3. (**F**) 4.1Å Cryo-EM structure of the Late LptD Intermediate. LptD and LptE are shown in orange and red, respectively. (**G**) Surface model of in-membrane cross section of the Late LptD Intermediate. The hybrid sheet and lateral seal are indicated. (**H**) Top-down overlay of fully folded LptDE (white, PDB: 4RHB) with the Late LptD Intermediate substrate (LptD and LptE in orange and red, respectively). Several extracellular loops have been hidden for clarity.

### Folding of LptD

To test whether our observations could be generalized to another substrate, we examined LptD, a substrate with many more β-strands than BamA, an unusual shape, and a soluble lipoprotein, LptE, around which it folds. We previously found that we can trap the native LptD *in vivo* at a late stage in folding, just as we have done with BamA^S^ (*37*). Moreover, deletion of an amino acid residue, D330, far from the N-terminal strand of LptD, slows folding further, causing LptD(ΔD330) to accumulate at this late intermediate (*37*). We therefore overexpressed and purified LptD(ΔD330) with I259C in β-strand 3 crosslinked to BamA^M^ (D512C) and determined its structure by cryo-EM to a final resolution of 4.1Å (Fig. 4F, Extended Data Fig. 10). Three features of the structures of BamA^S^ folding on BAM are also present in the structure of LptD folding on BAM. First, the machine and substrate form a hybrid sheet in which the open edge of the substrate faces the BamA^M^ lumen. Second, the C-terminal strand of the machine forms a sliding hydrophobic seal with the growing barrel (Fig. 4G). Third, the three N-terminal strands 1, 2, and 3 of the BamA^M^ machine barrel flex backward, with respect to superposed strands 4-6, at the late intermediate stage (compare with Fig 3C, right two panels). While the conformation of BamA^M^ is thus similar in the structures of the two late intermediates, the substrate conformations are not. BamA^S^ resembles the native BamA barrel and LptD resembles its native folded form, including LptE assembled within it (Fig. 4H) (*34–36*). We conclude that the BAM machine folds substrates of different sizes, shapes and compositions by the same general mechanism. Moreover, the features that determine barrel release are encoded in a capacity of the substrate to fold into a conformation that closely resembles its native structure, thereby bringing together the N- and C-terminal strands and enabling release by strand exchange.

## Discussion

BAM can fold a hundred different substrates which vary in size, shape, and composition. How it can do so has been a question since it was discovered. BAM has only two essential substrates: BamA, which catalyzes the folding process (*4*), and LptD, which together with LptE forms the outer membrane translocon that puts lipopolysaccharide in the outer leaflet of the outer membrane (*41*). The three intermediates, described here, of BamA^S^ folding on BAM represent different stages in the folding process; the structure of an LptDE intermediate folding on BAM, also described here, generalizes some of our conclusions. Several other groups have determined structures of the small barrel EspP on BAM (*18–20, 22*). Taken together, these structures show that the BamA machine barrel can distort at two regions to accommodate different stages of folding and different substrates (Fig. 5). As substrates fold progressively, they more and more resemble their final folded states, while the machine distorts to accommodate the maturing substrate barrel.

**Fig 5:**
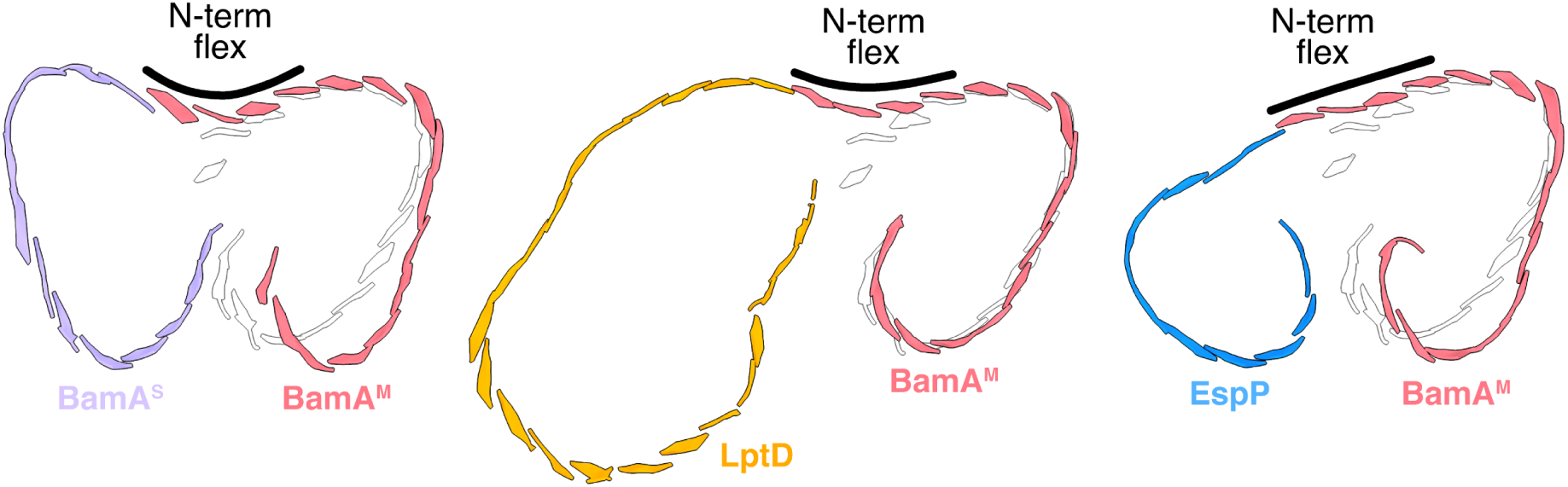
Distortions of BamA^M^ vary to accommodate its diverse substrates late in folding. In-membrane cross sections of late folding intermediates of BamA^S^ (left), LptD (middle), and EspP (right, PDB: 8BNZ). To visualize distortion present in the BamA^M^ barrel, BamA^M^ in each structure is aligned with and overlaid on apo BamA at β-strands 4-6 (light grey, PDB: 5LJO). Labels are present to illustrate the differential flexing of the N- terminal strands of BamA^M^ in each structure (black).

An important feature that BAM maintains during folding is the hydrophobic sliding seal between the outer surface of the most recently formed strands of the substrate barrel and the outer surface of the C-terminal region of the machine barrel. This seal, which is clearly visible in all the structures reported here, allows the unpaired H-bonding edge of the N-terminal strand to face into the lumen and template the next strand. In this way, the system avoids the potential sequestration of membrane lipids within the hybrid barrel lumen throughout folding. The hydrophobic interactions that maintain the seal allow it to slide as the substrate barrel grows until the N-terminal strand is close enough to the C-terminal strand that strand exchange can occur to cause release.

The studies here also show that small changes to substrates can slow the folding process. For example, a single amino acid deletion in strand 7 of LptD results in accumulation of a later, more N-terminal, strand within the BamA^M^ lumen (*37*). This functional LptD variant can complete folding, but at a slower rate than the wildtype variant (*37*, *38*). The structure of the Late LptD Intermediate containing a single amino acid deletion is very similar to the structure of the folded wildtype barrel. We have also shown that several BamA substrate variants fold more slowly on BAM than wildtype BamA. Thus, although BAM can support folding of a wide variety of substrates, specific features within individual substrates can affect the folding process. Structures of folding intermediates on BAM may be useful for guiding design of molecules that block late stages in the folding process.

## Methods

### SDS–PAGE and immunoblotting

Homemade Tris-HCl 4–20% polyacrylamide gradient gels were used with Tris-glycine running buffer. The 2× SDS sample loading buffer refers to a mixture containing 125 mM Tris (pH 6.8), 4% (w/v) SDS, 30% (v/v) glycerol, and 0.005% bromophenol blue. To analyze purified protein samples for cryo-EM, SDS–polyacrylamide gel electrophoresis (SDS–PAGE) was used followed by Coomassie staining with Coomassie blue (Alfa Aesar). 5 µL aliquots of purified protein samples were combined 1:1 with 2× SDS sample loading buffer with or without addition of 2% β-mercaptoethanol (BME). 10 µL of this mixture was loaded onto SDS-PAGE gels and run for 50 minutes at 200V, followed by Coomassie staining. Coomassie-stained SDS-PAGE gels were imaged using the gel setting on an Azure Biosystems C400 imager. For immunoblotting, SDS–PAGE gels were run for 90 min at 150 V. Proteins were transferred onto Immun-Blot PVDF membranes (Bio-Rad) and then incubated with the appropriate indicated antibodies. HRP conjugates were visualized with the Amersham ECL Prime western blotting detection kit (GE Healthcare). Immunoblots were imaged using the lowest sensitivity setting of the chemi feature of an Azure Biosystems C400 imager. An exposure time of 30 s was used. All immunoblots displayed are representative of three biological replicates. Full, uncropped immunoblot images can be found in Supplemental Figure 1.

### Plasmids, strains and oligonucleotides

Plasmids constructed for this study were made using components amplified by polymerase chain reaction (PCR) using KOD Hot Start DNA Polymerase (EMD Millipore). Gene fragments were then incubated with DpnI (New England Biolabs) and assembled via either Gibson assembly (*42*) (New England Biolabs) or circularization using kinase-ligase-DpnI (KLD) enzyme mix (New England Biolabs). Assembled plasmids were transformed into NovaBlue competent cells for replication. Plasmids, strains and oligonucleotides used in this study are reported in Supplemental Tables 1, 2, and 3 respectively.

### Expression and purification of disulfide-trapped BAM intermediate complexes for cryo-EM

Purification of disulfide-trapped BAM intermediate complexes was performed based on previous techniques (*17*). In particular, a BamA substrate (BamA^S^) containing a deletion of the periplasmic polypeptide-transport-associated (POTRA) domains 3-5, but no alterations to the barrel domain besides a single crosslinking cysteine, was used. The deletion of POTRAs 3-5 prevents BamA^S^ from binding lipoproteins and becoming an active machine (*43*) but does not affect the barrel’s ability to fold (*17*, *43*). For the Early Intermediate, the plasmid pJH114 (*44*) was modified by deleting the 6X-His tag on the C-terminus of BamE, inserting an 8X-His tag on the N-terminus of BamA, and introducing the D512C substitution, generating pBDT513. A BamA^S^ Early Intermediate overexpression plasmid was generated by deleting POTRA domains 3-5 from BamA, inserting a 2X-Strep tag on the N-terminus of BamA, introducing the Y637C substitution, and cloning the BamA^S^ construct into pBAD33, generating pBDT422. pBDT513 and pBDT422 were co-transformed into BL21 (DE3) cells and plated on selective Luria-Bertani agar (LB-agar) media containing 50 μg/ml carbenicillin and 25 μg/ml chloramphenicol. Clonal cells were diluted into 100mL LB containing 50 μg/ml carbenicillin and 25 μg/ml chloramphenicol and the resulting culture was grown to saturation overnight (37 °C, 200 rpm). The overnight culture was diluted 1:100 into 6×1.5 L flasks of LB supplemented with 50 μg/ml carbenicillin and 25 μg/ml chloramphenicol. The resulting cultures were grown (37 °C, 200 rpm) until an OD600 of approximately 0.7 was reached. At this point, protein expression was induced via addition of isopropyl β- d- 1-thiogalactopyranoside (VWR) and L-(+)-arabinose (Thermo Scientific) at final concentrations of 0.2 mM and 0.1% (w/v), respectively. Copper (II) (1,10-phenanthroline)_3_ was synthesized via addition of 45 mL of 30 mM copper (II) sulfate (Argos Organics) in water to 45 mL of 90 mM 1,10-phenanthroline (Sigma) in ethanol. After 30 minutes of culture shaking post-induction, oxidative disulfide crosslinking of BamA^M^ and BamA^S^ was induced via 1:100 addition of 30mM Copper (II) (1,10-phenanthroline)_3_ to expressing cultures. The cultures were allowed to continue shaking for an additional 2.5 hours. Cells were harvested via centrifugation (4200g, 15 minutes, 4 °C). Cell pellets were resuspended in a total of 300 mL PBS (20 mM NaH_2_PO_4_ pH 7.2, 150 mM NaCl) and centrifuged again (5,000g, 10 min, 4 °C). The resulting cell pellets were flash frozen in liquid nitrogen and stored at −80 °C before subsequent purification.

Cell pellets were thawed and resuspended in 150 mL buffer containing 20 mM Tris-HCl pH 8.0, 150 mM NaCl, 100 μg/ml lysozyme (Sigma-Aldrich), 1 mM PMSF (Sigma-Aldrich), and 25 μg/ml DNase I (Sigma-Aldrich). Cells were lysed using an Emulsiflex C5 (Avestin) at a pressure of 10,000 to 15,000 psi. After lysis, cell debris was removed via centrifugation (5,000g, 10 min, 4 °C). Membrane fractions were isolated via ultracentrifugation using a 45 Ti rotor (Beckman Coulter) (37,000 rpm, 30 min, 4 °C) and an Optima XE-90 ultracentrifuge (Beckman Coulter). The membrane pellets were resuspended in 120 mL buffer containing 20 mM Tris-HCl pH 8.0, 150 mM NaCl, 100 μg/ml lysozyme and 1 mM PMSF. Membrane fractions were solubilized via overnight incubation with 0.75% n-dodecyl-β-D-maltopyranoside (DDM) (Anatrace) and 0.5% glyco-diosgenin (GDN) (Anatrace) on a rocking platform (4 °C). Unsolubilized material was removed via ultracentrifugation in a 70 Ti rotor (Beckman Coulter) (37,000 rpm, 30 min, 4 °C).

The resulting supernatant was removed and incubated with 6 mL Ni-nitrilotriacetic acid (NTA) resin (Qiagen) that had been pre-washed with 10 column volumes of buffer W1 (20 mM Tris-HCl pH 8.0, 150 mM NaCl, 10 mM imidazole pH 8.0, 0.02% GDN). After batch binding on a rocking platform (1 h, 4 °C), the resin was washed with 30 column volumes buffer W1. The column was then capped, and the resin was resuspended in 5 column volumes buffer E1 (20 mM Tris-HCl pH 8.0, 150 mM NaCl, 200 mM imidazole pH 8.0, 0.02% GDN). The resin was allowed to settle for 10 minutes before elution from the column.

The eluate was immediately incubated with 1.5 mL Strep-Tactin XT 4Flow resin (IBA Lifesciences) that had been pre-washed with 10 column volumes buffer W2 (100 mM Tris-HCl pH 8.0, 150 mM NaCl, 0.02% GDN). After batch binding on a rocking platform (1 h, 4 °C), the resin was washed with 15 column volumes buffer W2. The column was then capped, and the resin was resuspended in 10 column volumes buffer E2 (100 mM Tris-HCl pH 8.0, 150 mM NaCl, 1 mM EDTA, 50 mM D-biotin, 0.02% GDN). The resin was allowed to settle for 10 minutes before elution from the column.

The strep resin eluate was concentrated to approximately 1 mL using an Amicon Ultra 4-ml 100-kDa molecular-weight cutoff centrifugal concentrator (EMD Millipore). The sample was then applied to an ÄKTA Pure (GE Healthcare Life Sciences) for purification via size-exclusion chromatography using a Superdex 200 Increase 10/300 GL column. The protein was eluted in buffer containing 20 mM Tris-HCl pH 8.0, 150 mM NaCl and 0.004% GDN. After elution, protein corresponding to the center peaks of the chromatogram was concentrated to 1 mg/mL using an Amicon Ultra 4-ml 100-kDa molecular-weight cutoff centrifugal concentrator (EMD Millipore), followed by an Amicon Ultra 0.5-ml 100-kDa molecular-weight cutoff centrifugal concentrator (EMD Millipore). A final yield of approximately 15 μg of protein per liter of bacterial culture was obtained. Presence of a disulfide crosslinked species of sufficient purity to structurally characterize was determined via SDS-PAGE analysis in the presence or absence of β-mercaptoethanol (BME).

The Middle Intermediate was expressed and purified in the same manner as the Early Intermediate with small differences. The BamA^M^ overexpression plasmid pBDT382 was used. pBDT382 is identical to pBDT513 except it contains a C-terminal 6X-His tag on BamE rather than an N-terminal 8X-His tag on BamA^M^. The BamA^S^ Middle Intermediate overexpression plasmid pBDT393, which contains the G597C substitution on BamA^S^ β-strand 9, was used in place of pBDT482. Additionally, after growing cultures reached an OD600 of 0.7, the temperature was reduced to 30 °C, and cells were allowed to continue shaking for 20 minutes. Expression was then induced, and cells were allowed to shake for three hours free of (1,10-phenanthroline)_3_. Oxidative disulfide formation was induced via addition of 0.3 mM (1,10-phenanthroline)_3_ to cells after harvest via centrifugation (5,000 g, 10 min, 4 °C), and the resulting slurry was allowed to rock (30 min, 25 °C) before further centrifugation and freezing. A final yield of approximately 20 μg of Middle Intermediate protein per liter of bacterial culture was obtained.

The Late Intermediate was expressed and purified in the same manner as the Early Intermediate, except the BamA^S^ Late Intermediate overexpression plasmid pBDT517, which contains the I458C substitution in β-strand 3 of BamA^S^, was used in place of pBDT482. A final yield of approximately 20 μg of Late Intermediate protein complex per liter of bacterial culture was obtained.

The Late LptD Intermediate was expressed and purified similarly to the Early Intermediate with the following alterations. The Late LptD Intermediate substrate overexpression plasmid pTS1293 was used in place of pBDT482. pTS1293 consists of a pCDFDuet backbone containing LptE and LptD-ΔD330 with the I259C substitution in β-strand 3. BL21 (DE3) cells were transformed and grown in the presence of 50 μg/ml of spectinomycin in place of chloramphenicol. Saturated overnight culture was diluted 1:40 into overexpression cultures. After growing cultures reached an OD600 of 0.7, the temperature was reduced to 30 °C, and cells were allowed to continue shaking for 20 minutes before harvest. Cell membrane fractions were solubilized via incubation with 1% GDN rather than a mixture of GDN and DDM. Affinity chromatography batch binding was performed overnight rather than for one hour. After elution, protein corresponding to the center peaks of the chromatogram was concentrated to 1.5 mg/mL. A final yield of approximately 50 μg of protein per liter of bacterial culture was obtained.

### Preparation of cryo-EM grids and collection of cryo-EM data

For the Early Intermediate, purified protein complex was generated as described in the prior section. Quantifoil R 2/1 holey carbon 400-mesh copper grids (Electron Microscopy Sciences) were glow-discharged using a PELCO easiGlow Glow Discharge Cleaning System set to a pressure of 0.39 mBar, current of 15 mA, hold of 10 s and glow of 30 s. 4 µL of purified protein complex at a concentration of 1 mg/mL was applied to the glow-discharged grids. Grids were blotted and vitrified in liquid nitrogen cooled ethane using a Thermo Fisher Scientific Vitrobot Mark IV (Thermo Fisher Scientific) set to a temperature of 4 °C, humidity of 100 %, a blot time of 7 s, wait time of 0 s, and a blot force of 14. The grid was clipped and loaded onto a Titan Krios G3i electron cryo-microscope (Thermo Fisher) operated at 300 kV accelerating voltage. Image stacks (movies) were recorded on a Gatan Bioquantum K3 Imaging Filter (Gatan) in counting mode. A nominal magnification of 105,000 x and a pixel size of 0.83 Å was used. The slit of the energy filter was set to 20 eV. A 50 μm C2 aperture and 100 μm objective aperture were used. The electron dose rate was 18.5 e-per physical pixel per second, and the subframe time was 0.045 s. A total exposure time of 1.9 s resulted in 42 total frames and a total dose of 51.1 e^-^ per Å^2^ (about 1.2 e^-^ per Å^2^ per subframe). The multishot scheme in SerialEM (*45*) version 4.1 was used for automated data collection with settings of nine holes per stage move and five movies per hole. A target defocus range of −1.2 µm to −2.6 µm was used with a step size of 0.1 µm. Data collection was performed over the course of approximately 72 hours and 20,970 raw movies were collected.

For the Middle Intermediate, UltrAufoil R 1.2/1.3 300-mesh gold grids (Electron Microscopy Sciences) were glow-discharged in the same way as grids for the Early Intermediate. 3.5 µL of purified protein complex at a concentration of 1 mg/mL was applied to the glow-discharged grids. Grids were blotted, vitrified, and clipped as described for the Early Intermediate before loading onto a Titan Krios G3i electron cryo-microscope (Thermo Fisher) operated at 300 kV accelerating voltage. Movies were recorded on a Gatan Bioquantum K3 Imaging Filter (Gatan) in counting mode. The same magnification, pixel size, energy filter and aperture settings were used as for the Early Intermediate. The electron dose rate was 11.5 e-per physical pixel per second, and the subframe time was 0.065 s. A total exposure time of 3.2 s resulted in 49 total frames and a total dose of 53.0 e^-^ per Å^2^ (about 1.2 e^-^ per Å^2^ per subframe). The multishot scheme in SerialEM (*45*) version 4.1 was used for automated data collection with settings of nine holes per stage move and two movies per hole. A target defocus range of −1.0 µm to −2.4 µm was used with a step size of 0.1 µm. Data collection was performed over the course of approximately 72 hours and 18,537 raw movies were collected.

For the Late Intermediate, UltrAufoil R 1.2/1.3 300-mesh gold grids (Electron Microscopy Sciences) were glow-discharged as described for the Early Intermediate. 4.0 µL of purified protein complex at a concentration of 2.2 mg/mL was applied to the glow-discharged grids. Grids were blotted, vitrified, and clipped as described for the Early Intermediate before loading onto a Titan Krios G3i electron cryo-microscope (Thermo Fisher) operated at 300 kV accelerating voltage. Movies were recorded on a Gatan Bioquantum K3 Imaging Filter (Gatan) in counting mode. The same magnification, pixel size, energy filter and aperture settings were used as for the Early Intermediate. The electron dose rate was 13.7 e-per physical pixel per second, and the subframe time was 0.05 s. A total exposure time of 2.5 s resulted in 50 total frames and a total dose of 49.6 e^-^ per Å^2^ (about 1.0 e^-^ per Å^2^ per subframe). The multishot scheme in SerialEM (*45*) version 4.1 was used for automated data collection with settings of nine holes per stage move and two shots per hole. A target defocus range of −1.0 µm to −2.4 µm was used with a step size of 0.1 µm. Data collection was performed over the course of approximately 72 hours and 21,962 raw movies were collected.

For the Late LptD Intermediate, Quantifoil 2/1 400 mesh grids (Electron Microscopy Sciences) were glow-discharged as described for the Early Intermediate. 4.0 µL of purified protein complex at a concentration of 1.5 mg/mL was applied to the glow-discharged grids. Grids were blotted, vitrified, and clipped as described for the Early Intermediate before loading onto a Titan Krios G3i electron cryo-microscope (Thermo Fisher) operated at 300 kV accelerating voltage. Movies were recorded on a Falcon 4i camera (Thermo Fisher) using EPU (Thermo Fisher) with a nominal magnification of 165,000 x and a pixel size of 0.736 Å. Two datasets were collected. For the first dataset, the electron dose rate was 10.0 e^-^ per physical pixel per second, and the subframe time was 0.06 s. A total exposure time of 2.69 s resulted in 46 total frames and a total dose of 49.55 e^-^ per Å^2^ (about 1.1 e^-^ per Å^2^ per subframe). A target defocus range of −0.6 µm to - 2.0 µm was used. Data collection was performed over the course of approximately 24 hours and 12,437 raw movies were collected. For the second dataset, the electron dose rate was 11.79 e^-^ per physical pixel per second, and the subframe time was 0.029 s. A total exposure time of 2.4 s resulted in 86 total frames and a total dose of 52.24 e^-^ per Å^2^ (about 0.64 e^-^ per Å^2^ per subframe). A target defocus range of −0.6 µm to −2.0 µm was used. Data collection was performed over the course of approximately 24 hours and 9,344 raw movies were collected.

### Cryo-EM data processing and 3D volume reconstruction

Image processing steps were carried out within the cryoSPARC (*46*) software package (Structura Biotechnology). Structural biology applications used in this project were compiled and configured by SBGrid (*47*).

For the Early Intermediate, 20,970 raw movies were patch-motion corrected, dose-weighted and contrast-transfer function (CTF) corrected within cryoSPARC Live. The resulting micrographs were curated and 1,661 micrographs with outlying CTF fit resolution or outlying relative ice thickness were discarded. 2,252,778 particles were picked from the remaining 19,309 micrographs using a TOPAZ (*48*) picking algorithm trained using curated circular-blob-picked particles from a preliminary processing run.

These particles were extracted with Fourier cropping to half-resolution and subjected to one round of 2D classification in which 360,925 particles were discarded. Two rounds of 3D classification were performed on the remaining 1,891,853 particles using nested rounds of ab-initio 3D reconstruction into four volumes, and subsequent heterogeneous refinement among those volumes. Local motion correction was performed on the resulting 1,014,155 selected particles. The particles were re-extracted without Fourier cropping and subjected to an additional round of heterogeneous refinement. Non-uniform refinement was performed on 543,043 selected particles to a final resolution of 3.7Å. However, individual strands of the substrate barrel were not clearly resolved at this point. The particles were polished using reference-based motion correction and subjected to another round of heterogeneous refinement, followed by selection of 374,510 particles and non-uniform refinement. Focused 3D classification was then performed using a mask surrounding the BamA^M^-BamA^S^ hybrid barrel region of the density to better resolve the BamA^S^ substrate. 143,293 particles were selected. Non-uniform refinement was performed on these particles to a final resolution of 3.6Å. The raw map was sharpened using a B-factor of −100 and the sharpened map was used for model building and validation. Local resolution estimation for maps displayed in supplementary figures was performed in RELION (*49*). Maps for display in main text figures were sharpened using DeepEMhancer (*50*).

For the Middle Intermediate, 18,537 raw movies were patch-motion corrected, dose-weighted and CTF corrected. The resulting micrographs were curated and 1,991 micrographs with outlying CTF fit resolution were discarded. 2,884,911 particles were picked from the resulting 16,546 micrographs using a circular blob picker. These particles were extracted with Fourier cropping to half-resolution and subjected to one round of 2D classification in which 426,198 particles were discarded. The resulting 2,458,511 particles were used to generate four ab-initio reconstruction 3D volumes and then subjected to one round of heterogeneous refinement. Local motion correction was performed on 969,230 selected particles, and the particles were re-extracted without Fourier cropping. The particles were then subjected to three rounds of heterogeneous refinement and 683,863 particles were selected. Non-uniform refinement was performed on these particles to a resolution of 3.3Å, however individual strands of the BamA^S^ substrate barrel were not clearly resolved. Focused 3D classification was performed using a mask surrounding the BamA^M^-BamA^S^ hybrid barrel region. 418,088 particles were selected and polished using reference-based motion correction. The resulting 418,024 particles were subjected to a final round of focused 3D classification using a mask surrounding the BamA^M^-BamA^S^ hybrid barrel region. 208,738 selected particles were carried forward to non-uniform refinement to a final resolution of 3.3Å. The raw map was sharpened using a B-factor of −115 and the sharpened map was used for model building and validation. Local resolution estimation for maps displayed in supplementary figures was performed in RELION (*49*). Maps for display in main text figures were sharpened using DeepEMhancer (*50*).

For the Late Intermediate, 21,962 raw movies were patch-motion corrected, dose-weighted and CTF corrected. The resulting micrographs were curated and 2,265 micrographs with outlying CTF fit resolution or outlying relative ice thickness were discarded. 1,999,604 particles were picked from the remaining 19,697 micrographs using a TOPAZ (*48*) picking algorithm trained using curated template-picked particles from a preliminary processing run. These particles were extracted with Fourier cropping to half-resolution and subjected to one round of 2D classification in which 200,375 particles were discarded. The remaining 1,799,229 particles were subjected to two rounds of heterogeneous refinement using volumes generated from a preliminary processing run. 773,053 particles were selected. Local motion correction was performed on these particles and they were re-extracted with no Fourier cropping. An additional round of heterogeneous refinement using preliminary volumes was performed and 647,144 particles were selected. These particles were used to generate three ab-initio reconstruction 3D volumes and heterogeneous refinement was performed using those volumes. 406,587 particles were carried forward to non-uniform refinement to a final resolution of 3.9Å. However, substrate density was not clearly visible in the resulting volume. Two rounds of focused 3D classification were performed using a mask surrounding the BamA^M^-BamA^S^ hybrid barrel region. 130,151 particles were selected and subjected to a further four rounds of nested ab-initio reconstruction and heterogeneous refinement. 39,744 particles were selected and polished using reference-based motion correction followed by classification in a final round of nested ab-initio reconstruction and heterogeneous refinement. 36,580 particles were selected and used for non-uniform refinement to a final reconstruction volume of 3.9Å. To help resolve BamA^S^, local refinement was performed on the 36,580 particles using a mask surrounding the BamA^M^- BamA^S^ hybrid barrel to a resolution of 4.1Å. The resulting locally refined map of the barrel region and the prior non-uniform refinement map were individually sharpened using a B-factor of −141 and combined into a composite map for model building and validation using PHENIX (*51*). Local resolution estimation for maps displayed in supplementary figures was performed in RELION (*49*). The locally refined map and non-uniform refined map were individually sharpened using DeepEMhancer (*50*) and then combined into a composite map using PHENIX for display in the main text.

For the Late LptD Intermediate, 12,437 raw movies in the first dataset were patch-motion corrected, dose-weighted and CTF corrected. The resulting micrographs were curated and 1,485 micrographs with outlying CTF fit resolution or outlying relative ice thickness were discarded. 382,582 particles were picked from the remaining 10,952 micrographs using a TOPAZ (*48*) picking algorithm trained using curated blob-picked particles from a preliminary processing run. In the second dataset, 9,344 raw movies in the first dataset were patch-motion corrected, dose-weighted and CTF corrected. The resulting micrographs were curated and 528 micrographs with outlying CTF fit resolution or outlying relative ice thickness were discarded. 428,649 particles were picked from the remaining 8,816 micrographs using a TOPAZ (*48*) picking algorithm trained using curated blob-picked particles from a preliminary processing run. The particles were extracted with Fourier cropping to half-resolution and the particle stacks from each dataset were combined. The resulting 811,231 particles were subjected to one round of 2D classification in which 143,471 particles were discarded. The remaining 1,799,229 particles were subjected to three rounds of 3D classification via nested ab-initio reconstruction of three classes and heterogeneous refinement among those classes. 88,113 particles were selected. Non-uniform refinement was performed on the resulting particle stack to a final resolution of 4.5Å. Reference-based motion correction was performed on the particle stack and the resulting polished particles were extracted with no Fourier cropping. One round of 3D classification via ab-initio reconstruction followed by heterogeneous refinement was performed. 79,819 particles sorted into two classes were selected and subjected to non-uniform refinement which resulted in a final reconstruction resolution of 4.1Å. To help resolve LptD and LptE, local refinement was performed on these particles using a mask surrounding the BamA^M^-LptD hybrid barrel, including the LptE plug, to a resolution of 4.3Å. The resulting locally refined map of the barrel region and the prior non-uniform refinement map were individually sharpened using a B-factor of −112 and combined into a composite map for model building and validation using PHENIX (*51*). Local resolution estimation for maps displayed in supplementary figures was performed in RELION (*49*).

### Model building, refinement and validation

Atomic models were generated using previously structures of the BAM complex (PDB: 6V05 and 5LJO for the Early and Middle Intermediate structures, 5D0O and 5D0Q for the Late Intermediate structure) (*11*, *17*, *18*), as well as a model of LptD(ΔD330)-LptE generated using Alphafold multimer accessed through Colabfold (*52*, *53*). Models of polyalanine β-strands built into the extra strands of density present in the Early and Middle Intermediate maps were generated in ChimeraX (*54*). Model components were rigid-body docked into the sharpened cryo-EM density maps using ChimeraX. Machine and substrate barrels were then morphed into the densities using secondary structure restraints within ISOLDE (*55*), accessed through ChimeraX. Manual model building and inspection of residue-by-residue fit to the densities was then carried out in ISOLDE and WinCoot (*56*). The resulting models were then refined against the density maps using real space refinement through PHENIX (*51*) with secondary structure and reference model restraints active. Backbone models without side chains were built into the density for BamA^M^ POTRA domains 1 and 2, as the corresponding densities are only visible at a permissive contour level, and at low resolution. The final models were inspected for fit to the cryo-EM maps and model geometry was evaluated using MolProbity (*57*). Cryo-EM data collection statistics are summarized in Extended Data Table 1, as well as Extended Data Figures 2, 3, 4, and 10. Model validation statistics are detailed in Extended Data Table 1.

### BamA^M^-BamA^S^ disulfide crosslinking

Disulfide crosslinking was performed based on previous techniques (*17*, *37*). 6×His– BamA^M^ containing either BamA^M^-D512C or BamA^M^-I806C was cloned into the pZS21 vector to generate pBDT374 and pBDT507 respectively. 3×FLAG-tagged substrates (BamA^S^ containing a deletion of POTRA domains 3–5, a cysteine substitution, as well as a loop deletion in some cases) were cloned into the pTrc99a vector to generate pBDT376, pBDT378, pBDT500-503, pBDT510, pBDT511, pBDT514, pBDT523, pBDT534, pBDT536, pBDT538, pBDT553-554, pBDT572-574, pBDT584, and pBDT586-588.

MC4100 cells (*58*) were transformed with either pBDT505, pBDT374 or pBDT507 containing wild type BamA^M^, BamA^M^(D512C), or BamA^M^(I806C) respectively and plated on LB-agar media containing 50 μg/mL kanamycin. Resulting cells were then transformed with the appropriate substrate plasmid and plated on LB-agar containing 50 μg/mL kanamycin and 50 μg/mL carbenicillin. These cells were then grown overnight to saturation in LB supplemented with 50 μg/mL kanamycin, 50 μg/mL carbenicillin, and 0.1% (w/v) glucose (37 °C, 220 rpm). The next day, saturated cultures were diluted 1:100 into 30 mL of fresh LB containing 50 μg/mL kanamycin and 50 μg/mL carbenicillin with no glucose. The cultures were allowed to grow for approximately 5 hours until an OD600 of approximately 0.8 was reached (37 °C, 220 rpm). Each culture was normalized to an optical density of 0.7 and volume of 25 mL. Cells were harvested via centrifugation (5000g, 10 minutes, 4 °C). The supernatant was aspirated and the resulting cell pellet was frozen overnight at −80 °C. Cell pellets were then thawed and resuspended in 5 mL of buffer containing 20 mM Tris-HCl pH 8.0, 150 mM NaCl, 100 μg/ml lysozyme (Sigma-Aldrich), 1 mM PMSF (Sigma-Aldrich), 25 μg/ml DNase I (Sigma-Aldrich), 20mM imidazole, and 1% (w/v) Anzergent 3-14 (Anatrace). Cells were lysed via repeated flash-freeze-thawing in liquid nitrogen: tubes containing cell suspension were immersed in liquid nitrogen for 2 minutes, followed by immersion in room-temperature water for 15 minutes. This cycle was repeated thrice, after which the lysate was clear. At this point, 10 μL of lysate was removed and diluted in 40 μL of a 1:1 mixture of 2× SDS buffer and 1M Tris-HCl pH 8.0. The lysate samples were then boiled (10 min, 100 °C). 6 μL of lysate samples were run per lane on SDS-PAGE gels and immunoblotted as described above using α-FLAG-HRP conjugate antibodies to ensure even expression of BamA^S^ substrate in disulfide crosslinking experiments.

The remaining lysate was added to 200 μL NTA resin (Qiagen) that had been pre-washed with 10 resin volumes of buffer containing 20 mM Tris-HCl pH 8.0, 150 mM NaCl, 20 mM imidazole, and 0.1% (w/v) Anzergent 3-14 (Anatrace). After batch binding (1 h, 4 °C), the resin was washed three times with 5 mL of buffer containing 20 mM Tris-HCl pH 8.0, 150 mM NaCl, 20mM imidazole, and 0.1% (w/v) Anzergent 3-14 (Anatrace). Washing was performed by resuspension of resin in 5 mL of buffer followed by centrifugation (2000 g, 3 minutes, 4 °C) to compact the resin. After the third wash, His-tagged proteins and disulfide adducts were eluted from the resin via addition of 1.1 mL of buffer containing 20 mM Tris-HCl pH 8.0, 150 mM NaCl, 200 mM imidazole, and 0.1% (w/v) Anzergent 3-14 (Anatrace). The elution buffer was then concentrated by trichloroacetic acid (TCA) precipitation: 1 mL of elution buffer containing proteins was added to 100 μL of 100% (w/v) TCA in water. Samples were allowed to incubate (20 min, 4 °C) before centrifugation to pellet precipitated proteins (21000 g, 10 min, 4 °C). TCA-containing buffer was aspirated and the resulting protein pellets were solubilized in 26 μL of a 1:1 mixture 2× SDS buffer and 1M Tris-HCl pH 8.0. Samples were boiled (10 min, 100 °C), and then 6 μL of purified protein samples were run on SDS-PAGE gels and immunoblotted as described above using α-FLAG-HRP conjugate antibodies, and α-His-HRP conjugate antibodies. The α-His immunoblot was performed to ensure even expression and purification efficiency of His-tagged BamA^M^. The α-FLAG immunoblot was used to assess disulfide crosslinking efficiency.

## Data Availability

Electron microscopy maps have been deposited into the Electron Microscopy Data Bank (EMDB) under accession codes EMD-48253 (Early Intermediate), EMD-48254 (Middle Intermediate), EMD-48255 (Late Intermediate composite), EMD-48251 (Late Intermediate consensus), EMD-48252 (Late Intermediate focused refinement), EMD-71132 (Late LptD Intermediate consensus), EMD-71133 (Late LptD Intermediate focused refinement), EMD-71134 (Late LptD Intermediate composite). Atomic models have been deposited into the Protein Data Bank under accession codes 9MGE (Early Intermediate), 9MGF (Middle Intermediate), 9MGG (Late Intermediate), and 9P1U (Late LptD Intermediate). All other data are available in the article and the associated supplementary information.

## Acknowledgements

We thank R. Walsh, M. Mayer, C. Leistner, S. Huber, and S. Sterling for assistance with electron microscopy data collection; K. Pahil, M. Gilman and I. Shlosman for input on electron microscopy sample preparation; K. Pahil for input on electron microscopy data processing. B. Thomson thanks J. Hu. Cryo-EM data were collected at the Harvard Cryo-EM Center for Structural Biology at Harvard Medical School. Structural biology applications used in this project were compiled and configured by SBGrid. Funding for this work was provided by National Institutes of Health grants U19AI158028 and R01AI081059 to D.K. T.M.A.d.S was supported by a Howard Hughes Medical Institute Hanna H. Gray Postdoctoral Fellowship.

## Author contributions

Conceptualization: BDT, DK

Methodology: BDT, MDM, SR, TMAdS, DK

Investigation: BDT, MDM, TMAdS

Validation: BDT, SR

Visualization: BDT

Funding acquisition: DK

Project administration: DK

Supervision: DK

Writing – original draft: BDT, DK

Writing – review & editing: BDT, MDM, SR, TMAdS, SCH, DK

## Competing interests

The authors declare no competing interests.

## Additional information

Supplementary Information is available for this paper.

## Extended data tables

**Extended Data Table 1:** Cryo-EM data collection, refinement, and validation statistics.

|  | Early<br>Intermediate<br>(PDB 9MGE)<br>(EMD-48253) | Middle<br>Intermediate<br>(PDB 9MGF)<br>(EMD-48254) | Late<br>Intermediate<br>(PDB 9MGG)<br>(EMD-48255) | Late LptD<br>Intermediate<br>(PDB 9P1U)<br>(EMD-71132) |
| --- | --- | --- | --- | --- |
| **Data collection and processing** |  |  |  |  |
| Microscope | Krios | Krios | Krios | Krios |
| Magnification | 105,000 | 105,000 | 105,000 | 165,000 |
| Voltage (kV) | 300 | 300 | 300 | 300 |
| Electron exposure (e-/$\text{\AA}^2$) | 51.1 | 53 | 49.6 | 49.5/52.2 |
| Defocus range ($\mu\text{m}$) | 1.2-2.6 | 1.0-2.4 | 1.0-2.4 | 0.6-2.0 |
| Pixel size ($\text{\AA}$) | 0.83 | 0.83 | 0.83 | 0.74 |
| Symmetry imposed | C1 | C1 | C1 | C1 |
| Initial particle images (no.) | 2,252,778 | 2,884,911 | 1,999,604 | 811,231 |
| Final particle images (no.) | 143,293 | 208,738 | 39,744 | 79,819 |
| Global map resolution ($\text{\AA}$) | 3.6 | 3.3 | 3.9 | 4.1 |
| FSC threshold | 0.143 | 0.143 | 0.143 | 0.143 |
| **Refinement** |  |  |  |  |
| Initial model source | 5D0O, 5LJO | 6V05, 5LJO | 5D0O, 5LJO | 5LJO, AlphaFold |
| Model resolution ($\text{\AA}$) | 4.0 | 3.5 | 4.2 | 4.3 |
| FSC threshold | 0.5 | 0.5 | 0.5 | 0.5 |
| Map sharpening B-factor ($\text{\AA}^2$) | -100 | -115 | -141 | -112 |
| Model composition |  |  |  |  |
| Non-H atoms | 11740 | 12658 | 13296 | 15852 |
| Protein residues | 1581 | 1625 | 1774 | 2080 |
| Ligands | 0 | 0 | 0 | 0 |
| Model vs. data CC | 0.75 | 0.76 | 0.76 | 0.76 |
| R.m.s. deviations |  |  |  |  |
| Bond lengths ($\text{\AA}$) | 0.005 | 0.003 | 0.004 | 0.004 |
| Bond angles ($^\circ$) | 0.933 | 0.743 | 0.839 | 0.722 |
| Validation |  |  |  |  |
| Molprobrity score | 1.91 | 1.54 | 2.02 | 1.84 |
| Clashscore | 7.13 | 4.35 | 10.13 | 10.41 |
| Poor rotamers (%) | 1.10 | 0.37 | 1.03 | 0.25 |
| Ramachandran plot |  |  |  |  |
| Favored (%) | 92.16 | 95.25 | 92.11 | 95.61 |
| Allowed (%) | 7.84 | 4.62 | 7.83 | 4.20 |
| Disfavored (%) | 0.00 | 0.12 | 0.06 | 0.20 |

## Extended data figures

**Extended Data Fig. 1:**
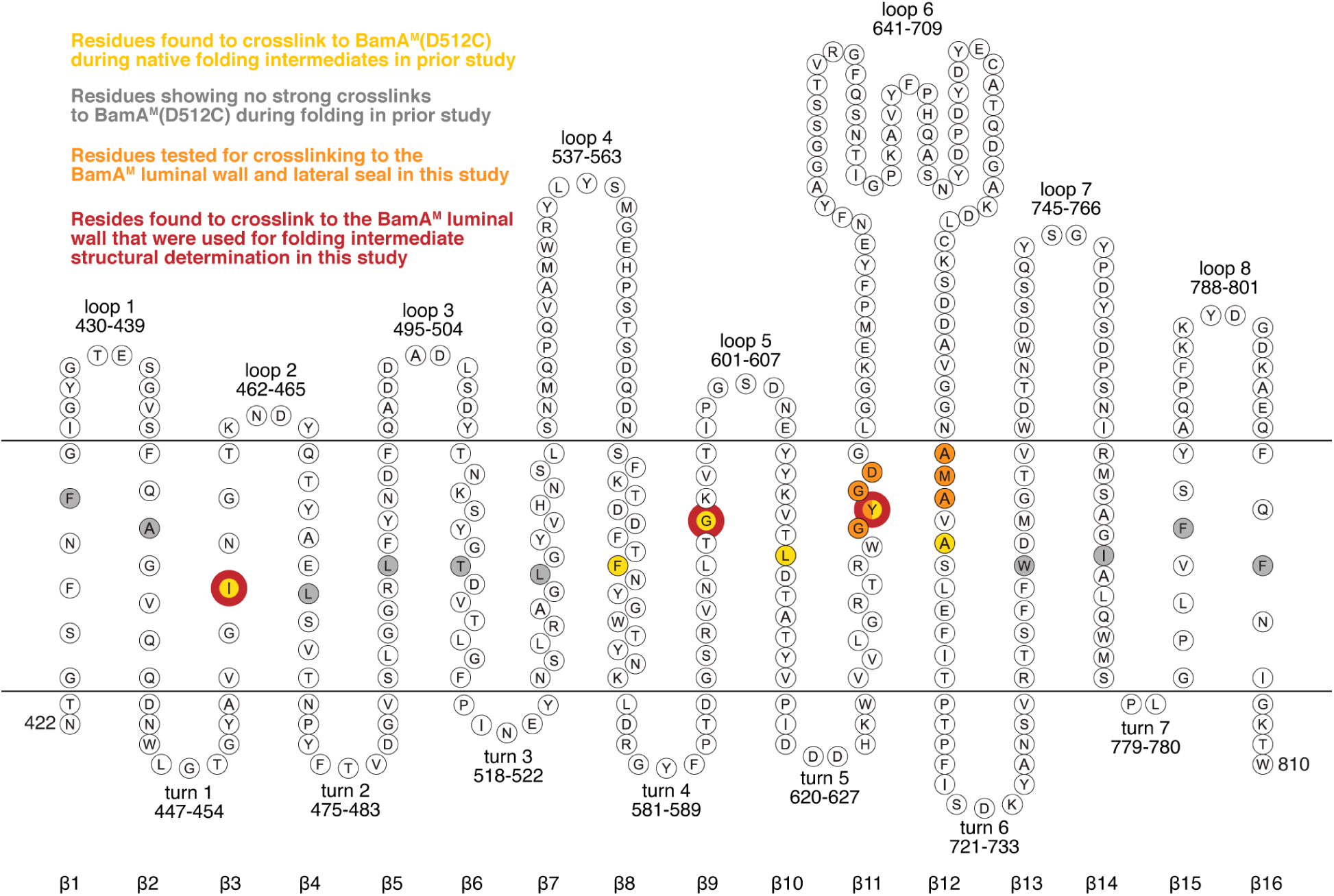
Topological depiction of the BamA β-barrel domain. The extracellular loops, periplasmic turns, and transmembrane β-strands are labeled. Residues within BamA^S^ β-strands that, upon substitution with cysteine, were previously found to crosslink to BamA^M^(D512C) during native folding intermediates are highlighted in yellow (*37*). Tested residues within β-strands that were found to show no such strong crosslinking are shown in gray. Residues that were substituted for cysteines and used in crosslinking experiments to the BamA^M^ luminal wall and to the BamA^M^ C-terminal region at the lateral seal are highlighted in orange. Residues that were substituted for cysteines to trap folding BamA^S^ within native folding intermediates for structural characterization are outlined with red circles. These substitutions were used to purify, from left to right, the Late Intermediate (BAM-BamA^S^ (I458C)), Middle Intermediate (BAM-BamA^S^ (G597C)), and Early Intermediate (BAM-BamA^S^ (Y637C)) complexes.

**Extended Data Fig. 2:**
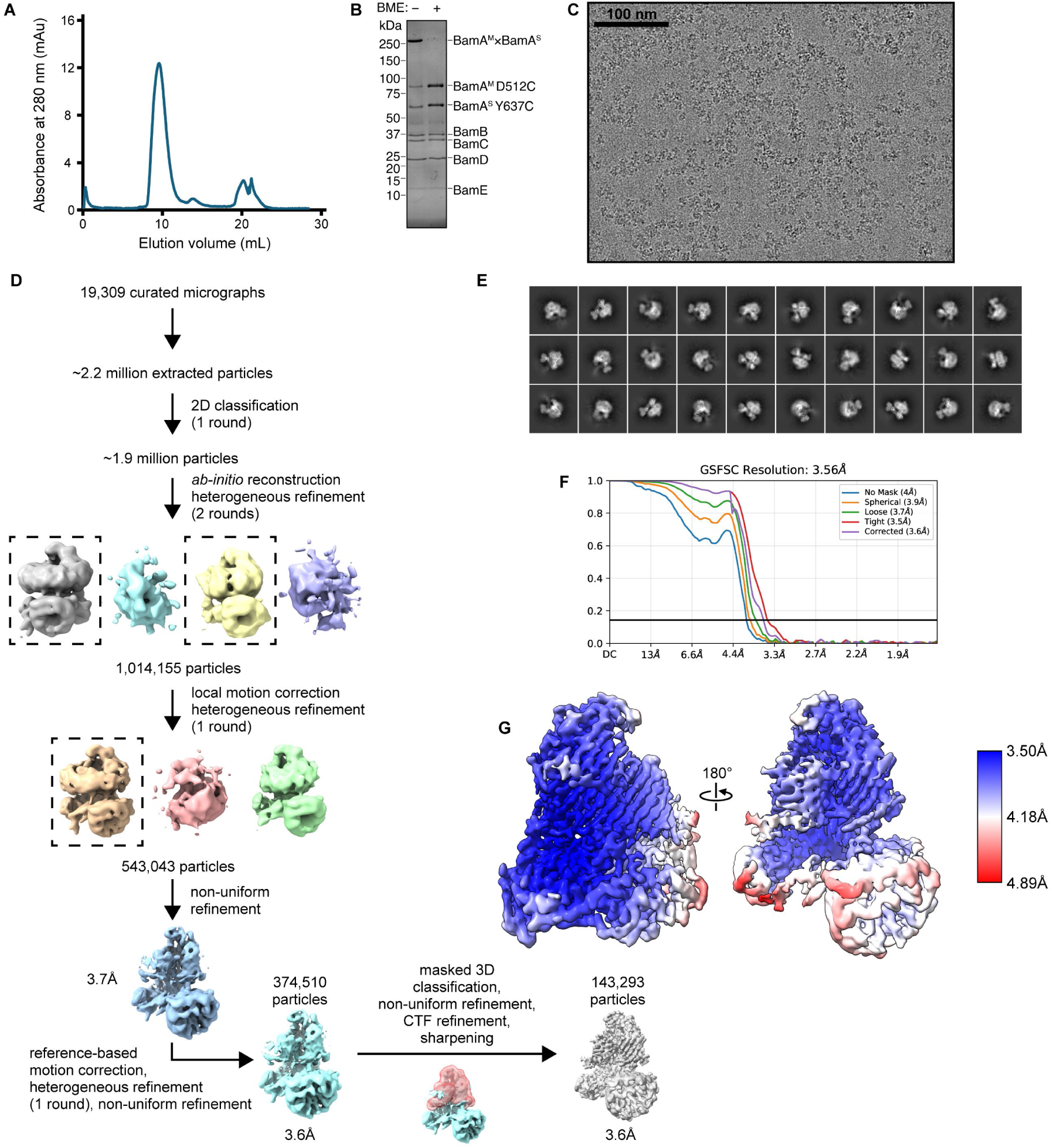
Cryo-EM data processing and analysis for the Early Intermediate complex. (**A**) Representative size-exclusion chromatogram of the Early Intermediate complex. The major peak fractions were collected and pooled before concentration and freezing for cryo-EM grid preparation. (**B**) Coomassie-stained SDS-PAGE gel of purified Early Intermediate complex. The high-molecular-weight band present within the sample reduces to BamA^M^ and BamA^S^ upon addition of β-mercaptoethanol (BME). (**C**) Representative cryo-EM micrograph of the Early Intermediate complex embedded in vitreous ice. (**D**) Scheme of cryo-EM data processing for the Early Intermediate complex. Images of cryo-EM densities were created using ChimeraX. Focused 3D classification used a mask that surrounded the hybrid barrel region of the density. (**E**) Representative 2D class averages of extracted Early Intermediate cryo-EM particles. (**F**), Gold-standard Fourier shell correlation (FSC) curves calculated with different masks within cryoSPARC. The resolution was determined at FSC = 0.143 (horizontal line). The final corrected mask gave a resolution of 3.6Å. (**G**) Final map of the Early Intermediate non-uniform refined density. The density has been colored by local resolution calculated using RELION. The map used for model building and validation was sharpened with a uniform B-factor of −100. Maps displayed in the main text figures were postprocessed with DeepEMhancer.

**Extended Data Fig. 3:**
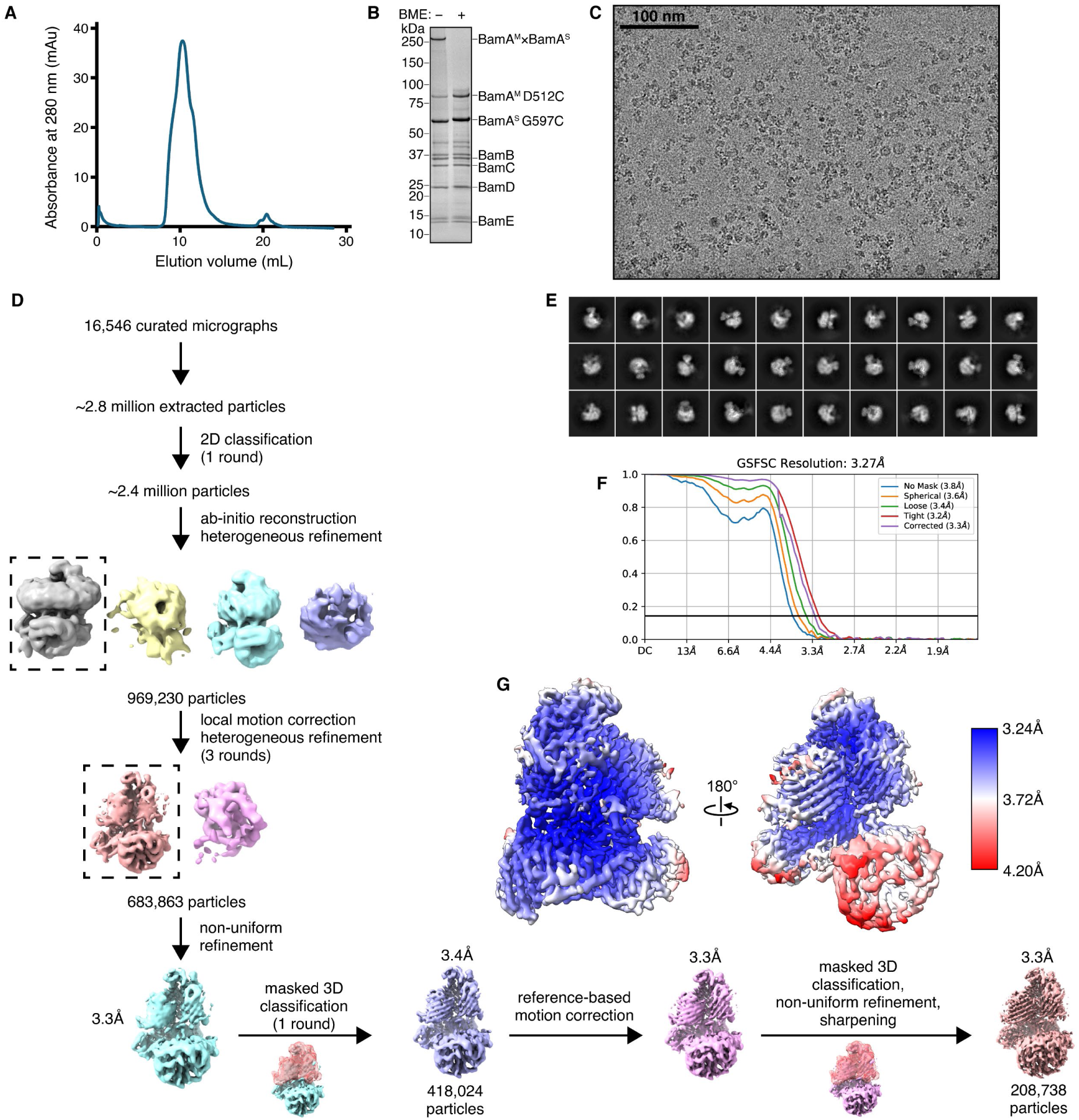
Cryo-EM data processing and analysis for the Middle Intermediate complex. (**A**) Representative size-exclusion chromatogram of the Middle Intermediate complex. The major peak fractions were collected and pooled before concentration and freezing for cryo-EM grid preparation. (**B**) Coomassie-stained SDS-PAGE gel of purified Middle Intermediate complex. The high-molecular-weight band present within the sample reduces to BamA^M^ and BamA^S^ upon addition of β-mercaptoethanol (BME). (**C**) Representative cryo-EM micrograph of the Middle Intermediate complex embedded in vitreous ice. (**D**) Scheme of cryo-EM data processing for the Middle Intermediate complex. Images of cryo-EM densities were created using ChimeraX. Focused 3D classification used a mask that surrounded the hybrid barrel region of the density. (**E**) Representative 2D class averages of extracted Middle Intermediate cryo-EM particles. (**F**) Gold-standard Fourier shell correlation (FSC) curves calculated with different masks within cryoSPARC. The resolution was determined at FSC = 0.143 (horizontal line). The final corrected mask gave a resolution of 3.3Å. (**G**) Final map of the Middle Intermediate non-uniform refined density. The density has been colored by local resolution calculated using RELION. The map used for model building and validation was sharpened with a uniform B-factor of −115. Maps displayed in the main text figures were postprocessed with DeepEMhancer.

**Extended Data Fig. 4:**
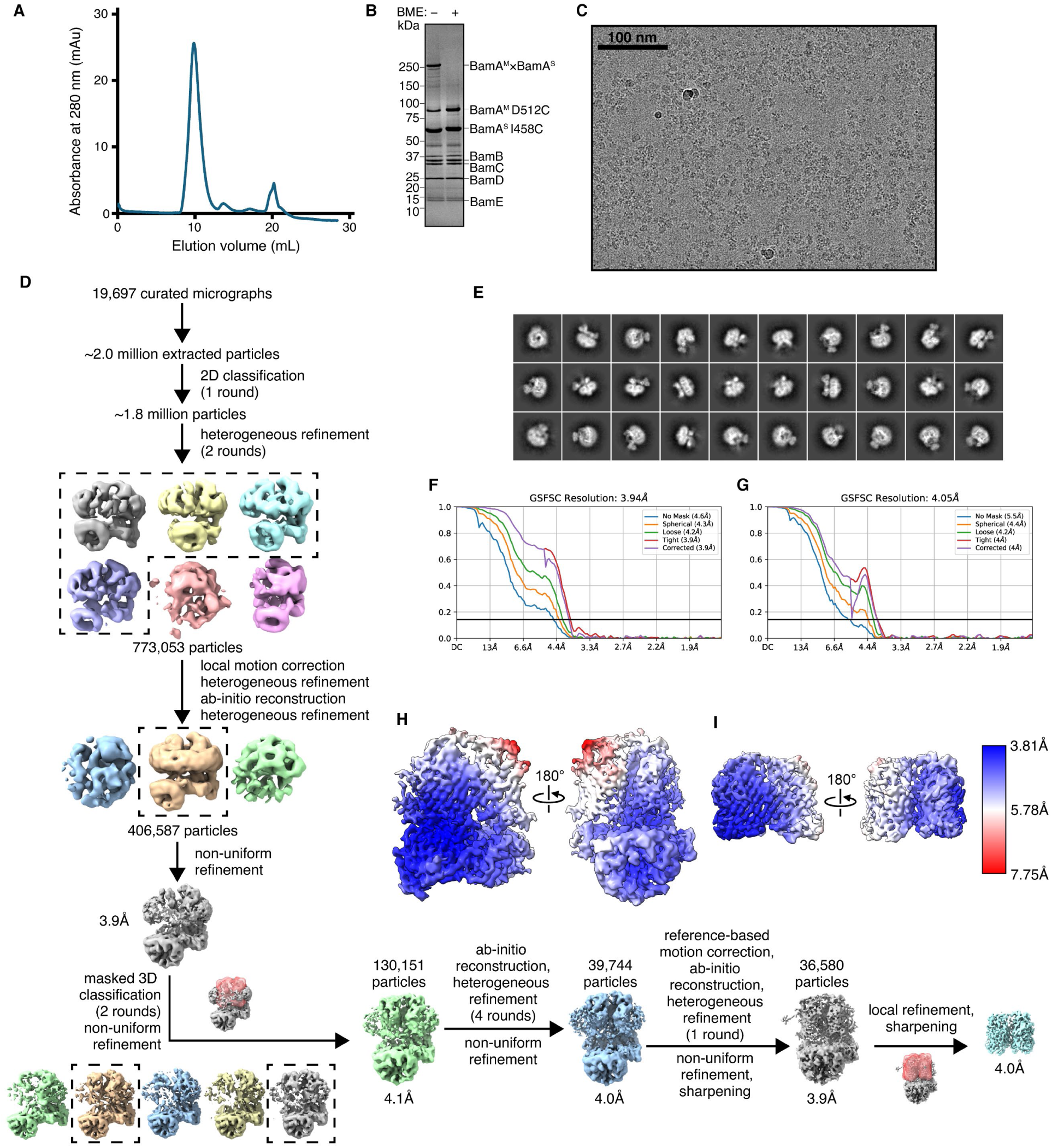
Cryo-EM data processing and analysis for the Late Intermediate complex. (**A**) Representative size-exclusion chromatogram of the Late Intermediate complex. The major peak fractions were collected and pooled before concentration and freezing for cryo-EM grid preparation. (**B**) Coomassie-stained SDS-PAGE gel of purified Late Intermediate complex. The high-molecular-weight band present within the sample reduces to BamA^M^ and BamA^S^ upon addition of β-mercaptoethanol (BME). (**C**) Representative cryo-EM micrograph of the Late Intermediate complex embedded in vitreous ice. (**D**) Scheme of cryo-EM data processing for the Late Intermediate complex. Images of cryo-EM densities were created using ChimeraX. Focused 3D classification and local refinement masks surrounded the hybrid barrel region of the density. (**E**) Representative 2D class averages of extracted Late Intermediate cryo-EM particles. (**F, G**) Gold-standard Fourier shell correlation (FSC) curves calculated with different masks within cryoSPARC from the global non-uniform refined (F) and locally refined (G) densities of the Late Intermediate complex. The resolution was determined at FSC = 0.143 (horizontal line). The final corrected mask gave a resolution of 3.9Å for the global non-uniform refined density, and a resolution of 4.1Å for the locally refined density. (**H, I**) Final maps of the Late Intermediate non-uniform refined density (H) and locally refined density (I). The density has been colored by local resolution calculated using RELION. Final maps used for model building were individually sharpened with a uniform B-factor of −141 and combined into a composite map using PHENIX. Maps displayed in the main text figures were postprocessed with DeepEMhancer and then combined using PHENIX.

**Extended Data Fig. 5:**
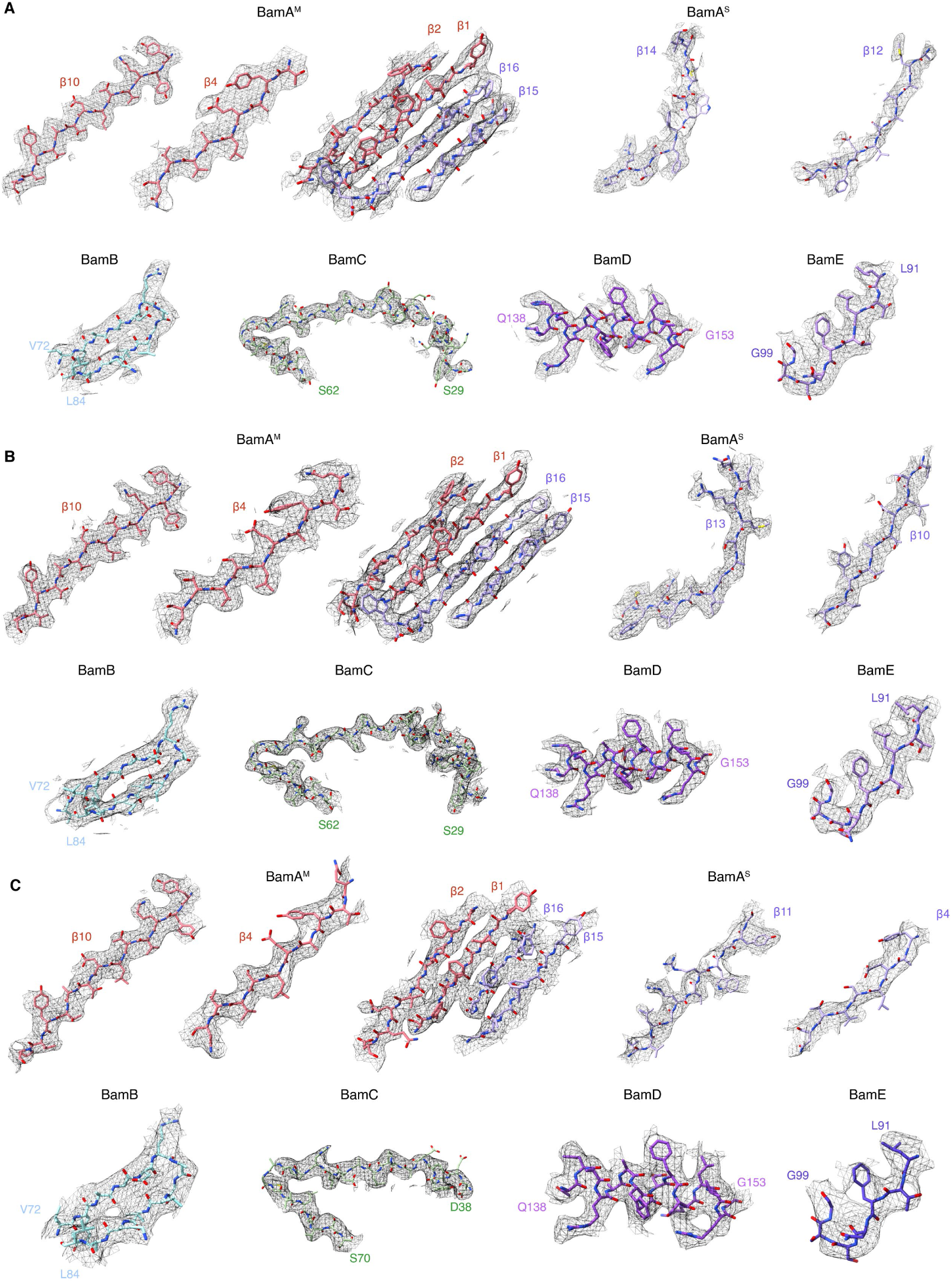
Fit of the atomic models into the BamA^S^ intermediate cryo-EM maps. (**A-C**) Atomic models and their fit into the sharpened cryo-EM maps for selected parts of each chain in the Early Intermediate (A), the Middle Intermediate (B), and the Late Intermediate (C) structures. Images were generated in ChimeraX using a carve radius of 2.5Å.

**Extended Data Fig. 6:**
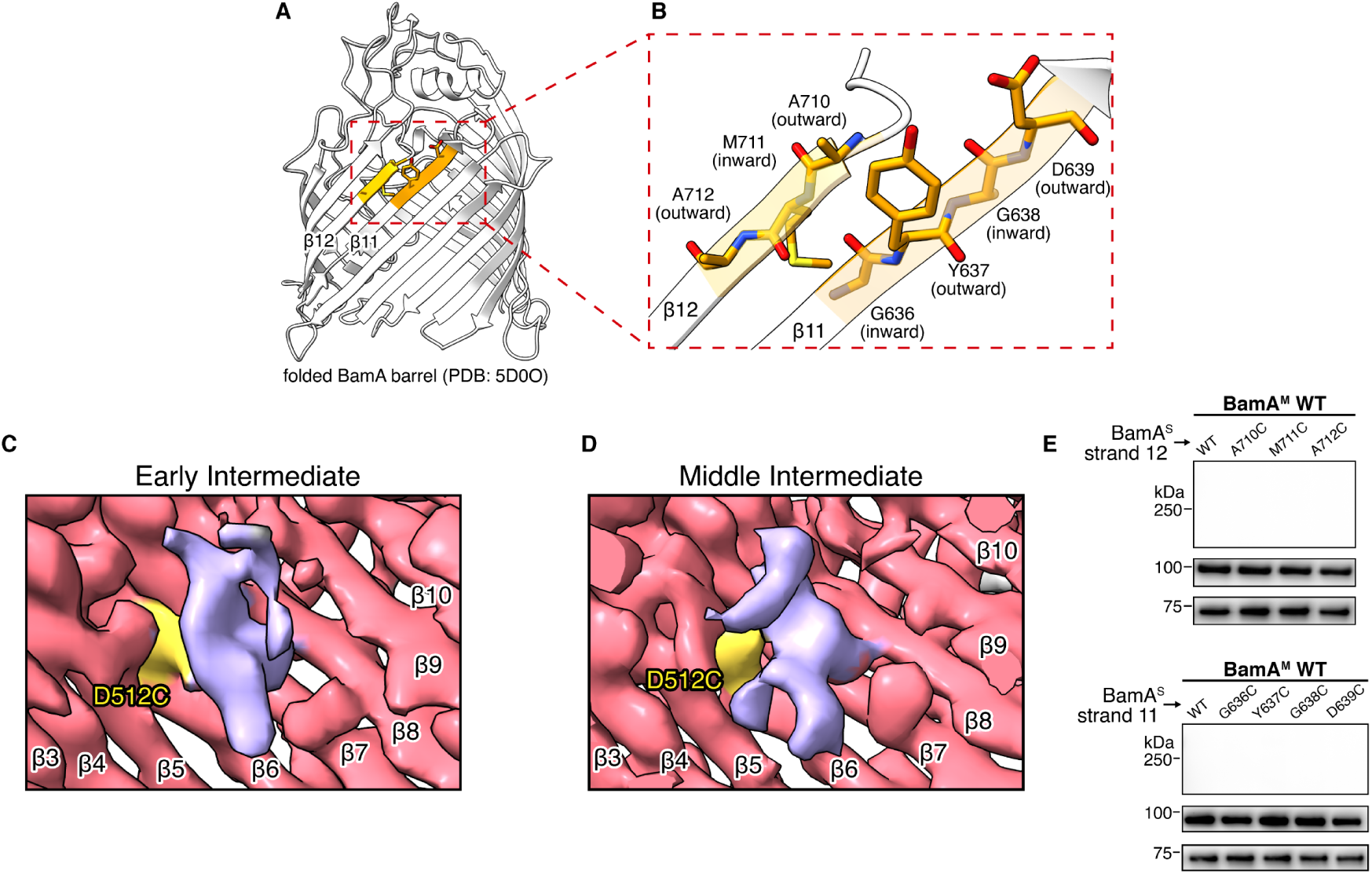
BamA^S^ density within the BamA^M^ barrel. (**A**), Locations of β-strands 11 and 12 within the closed, folded BamA barrel structure. The residues within β-strands 11 and 12 that were substituted for cysteines and used for disulfide crosslinking experiments to the BamA^M^ luminal wall and to the BamA^M^ C-terminal region at the lateral seal are highlighted in orange. (**B**), Region of the folded BamA barrel that contains the residues tested for luminal wall and lateral seal disulfide crosslinking. When organized into the native sheet, A710 and A712 within β-strand 12 have outward-facing side chains pointed toward the membrane while M711 has an inward-facing side chain pointed toward the barrel lumen. Similarly, when organized into the native sheet, Y637 and D639 have outward-facing side chains while G636 and G638 would have inward-facing side chains upon substitution with cysteine. (**C**, **D**) Inside-the-barrel view of the BamA^M^ luminal wall for the Early Intermediate (C) and Middle Intermediate (D) cryo-EM densities. BamA^M^ strands are labeled. Both maps contain density within the BamA^M^ barrel at the density at the point of crosslinking between the machine and substrate: BamA^M^(D512C) (yellow). We interpret this density as corresponding to the disordered region of the substrate containing the crosslinking substrate cysteine in each complex (β-strand 11 for the Early Intermediate, and β-strand 9 for the Middle Intermediate). (**E**) Expression of BamA^S^ cysteine substitutions in β-strand 12 (upper) and β-strand 11 (lower) alongside wild-type BamA^M^ containing no cysteine substitution results in no disulfide crosslinking bands. Disulfide crosslinks between BamA^M^(D512C) or BamA^M^(I806C) and BamA^S^ cysteine-containing variants are therefore dependent upon the engineered BamA^M^ cysteine. Data are laid out as described in Fig. 2D.

**Extended Data Fig. 7:**
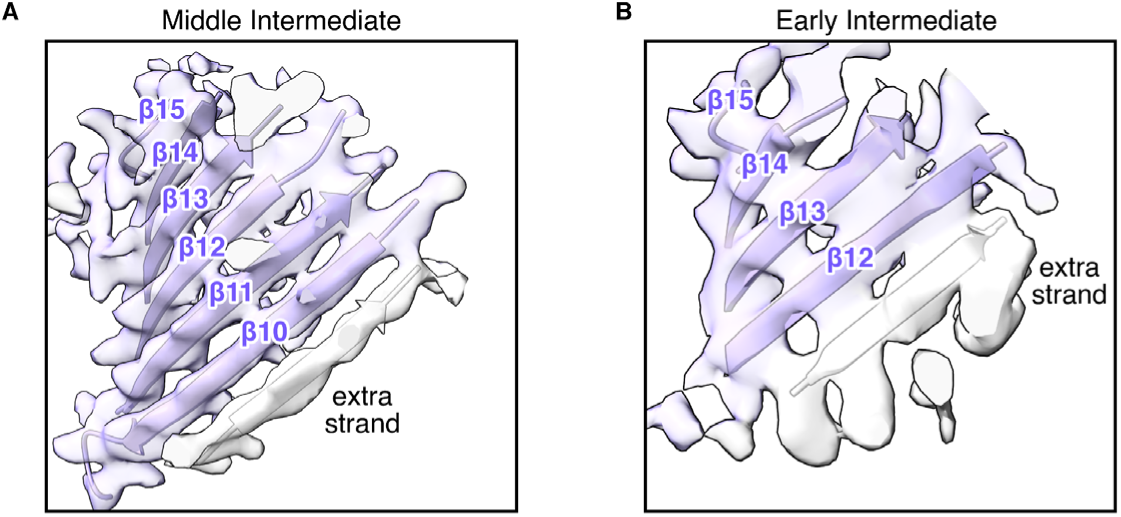
Substrate density in the Early and Middle Intermediate structures. (**A**, **B**) Detail view of the BamA^S^ substrate density at the interface of the substrate edge and the BamA^M^ lumen for the Middle Intermediate (A) and Early Intermediate (B). An “extra strand” of density is visible for each intermediate but not featured enough to clearly identify (light grey). A polyalanine β-strand sequence was built into this density in the Early and Middle Intermediate models.

**Extended Data Fig. 8:**
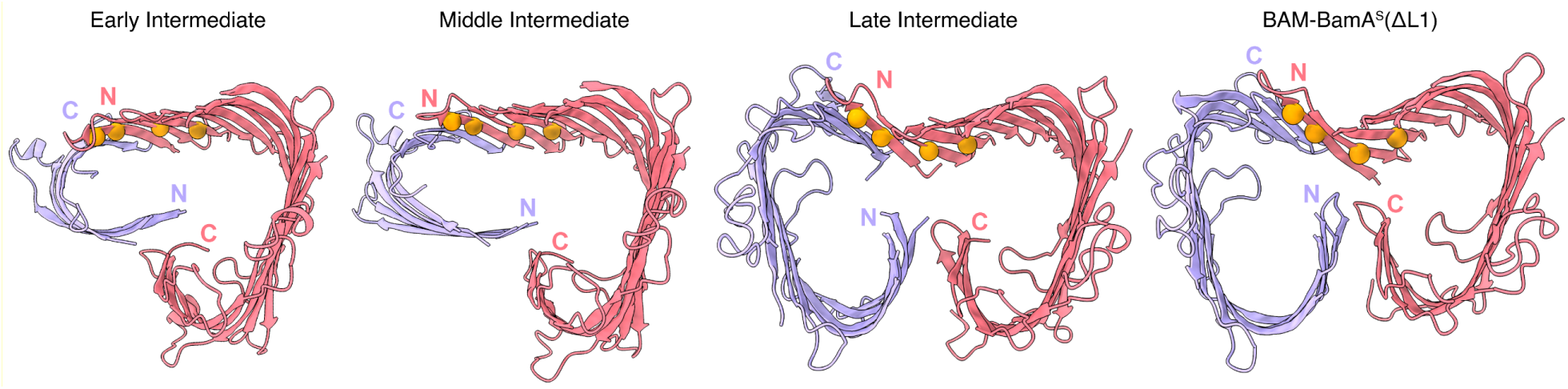
The region of BamA^M^ that flexes outward contains several inward-oriented glycines. Top-down depictions of the hybrid barrel domains of, from left to right, the Early Intermediate, Middle Intermediate, Late Intermediate, and BAM-BamA^S^(ΔL1). Four inward-oriented glycine residues within BamA^M^ β-strands 1-3, the machine’s N-terminal β-strands that progressively flex outward during folding, are indicated (orange spheres). Several extracellular loops have been hidden for clarity.

**Extended Data Fig. 9:**
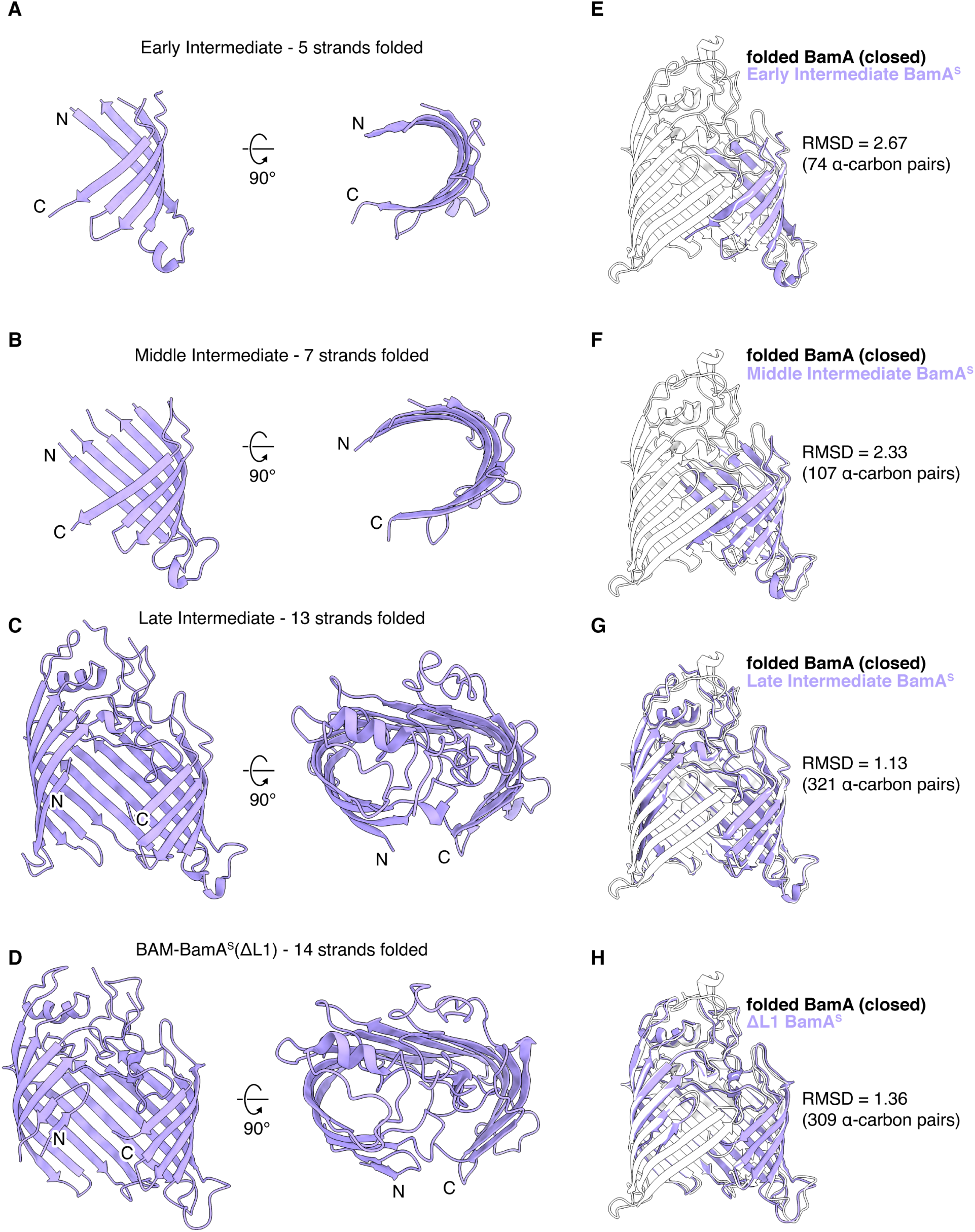
Folded portions of the BamA^S^ substrate barrel in progressive folding intermediates. (**A**-**D**), Side view (left) and top-down view (right) of the BamA^S^ substrate barrel within the Early Intermediate (A), Middle Intermediate (B), Late Intermediate (C) and BAM-BamA^S^(ΔL1) (D) structures. As strands are added, the substrate’s open N-terminal edge approaches the C-terminal strand. (**E**-**H**) Overlays of the fully folded closed BamA barrel (white, PDB: 5D0O) and the BamA^S^ substrate (lavender) for the Early Intermediate (E), Middle Intermediate (F), Late Intermediate (G) and BAM-BamA^S^(ΔL1) (H). Overall, the substrate’s folded strands closely resemble the corresponding portion of the folded, closed BamA^S^ barrel throughout folding. PDB: 5D0O) and the BamA^S^ substrate (lavender) for the Early Intermediate (E), Middle Intermediate (F), Late Intermediate (G) and BAM-BamA^S^(ΔL1) (H). Overall, the substrate’s folded strands closely resemble the corresponding portion of the folded, closed BamA^S^ barrel throughout folding.

**Extended Data Fig. 10:**
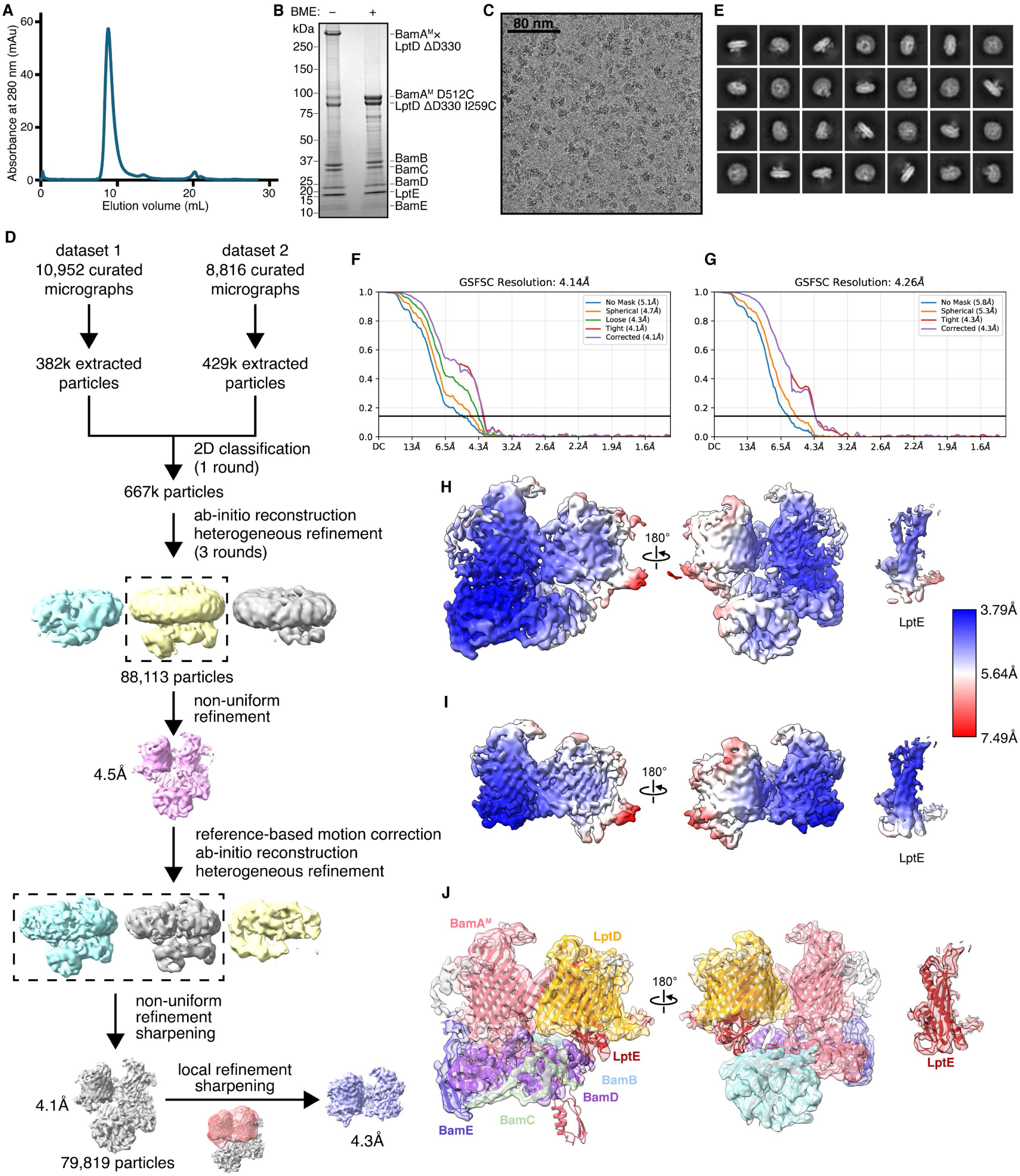
Cryo-EM data processing and analysis for the Late LptD Intermediate complex. (**A**) Representative size-exclusion chromatogram of the Late LptD Intermediate complex. The major peak fractions were collected and pooled before concentration and freezing for cryo-EM grid preparation. (**B**) Coomassie-stained SDS-PAGE gel of purified Late LptD Intermediate complex. The high-molecular-weight band present within the sample reduces to BamA^M^ and LptD upon addition of β-mercaptoethanol (BME). (**C**) Representative cryo-EM micrograph of the Late LptD Intermediate complex embedded in vitreous ice. (**D**) Scheme of cryo-EM data processing for the Late LptD Intermediate complex. Images of cryo-EM densities were created using ChimeraX. The local refinement mask surrounded the hybrid barrel region of the density, including the LptE plug. (**E**) Representative 2D class averages of extracted Late Intermediate cryo-EM particles. (**F**, **G**) Gold-standard Fourier shell correlation (FSC) curves calculated with different masks within cryoSPARC from the global non-uniform refined (F) and locally refined (G) densities of the Late Intermediate complex. The resolution was determined at FSC = 0.143 (horizontal line). The final corrected mask gave a resolution of 4.1Å for the global non-uniform refined density, and a resolution of 4.3Å for the locally refined density. (**H**, **I**) Final maps of the Late LptD Intermediate non-uniform refined density (H) and locally refined density (I). The density has been colored by local resolution calculated within RELION. Final maps used for model building were individually sharpened with a uniform B-factor of −112 and combined into a composite map using PHENIX. Density for the LptE plug produced using a carve radius of 2.5Å around LptE is shown (right). (**J**) Fit of atomic model into the Late LptD Intermediate cryo-EM map. Models and densities corresponding to BamA^M^, BamB, BamC, BamD, BamE, LptD, and LptE are colored in salmon, light blue, light green, purple, dark blue, orange, and red respectively. The composite map of local refined and global refined density is shown. Density for the LptE plug produced using a carve radius of 2.5Å around LptE is shown (right).

## Notes

### Competing Interest Statement

The authors have declared no competing interest.

