## Supplementary Information for "Structures of folding intermediates on BAM show diverse substrates fold by a uniform mechanism"

**Supplementary Figure 1: Uncropped gels and blots for data shown in figures.** For each figure, the cropped regions shown are denoted by boxes. Immunoblots shown are representative of three biological replicates.

Figure 2D

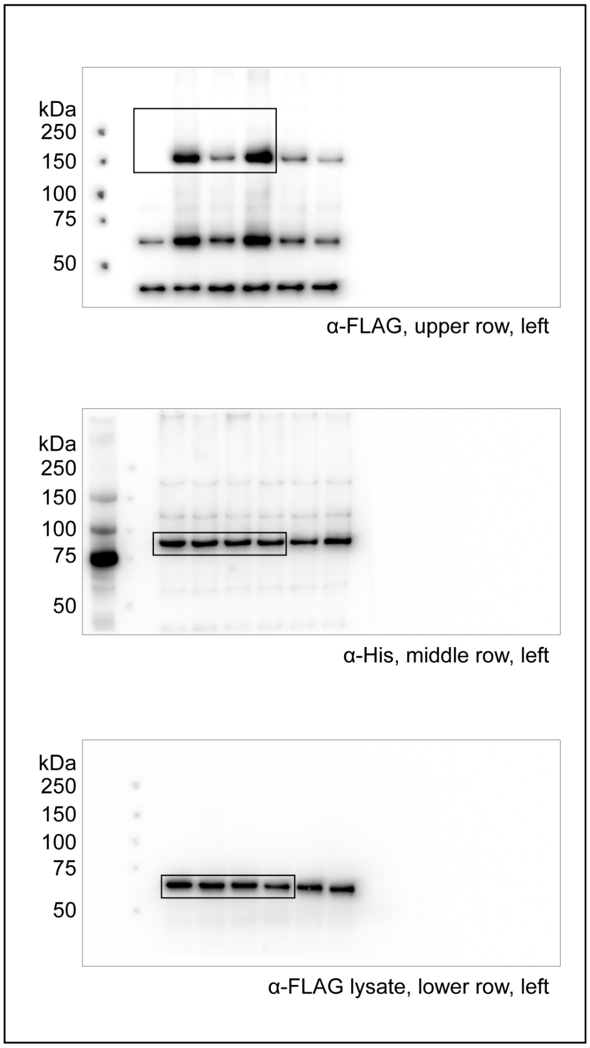


Figure 2I

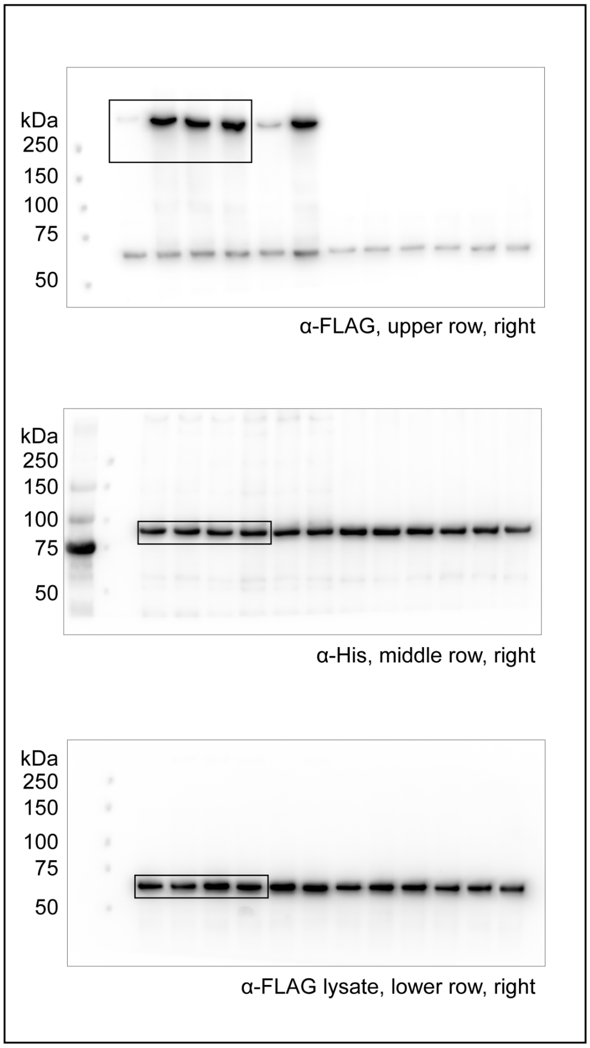

Figure 2H

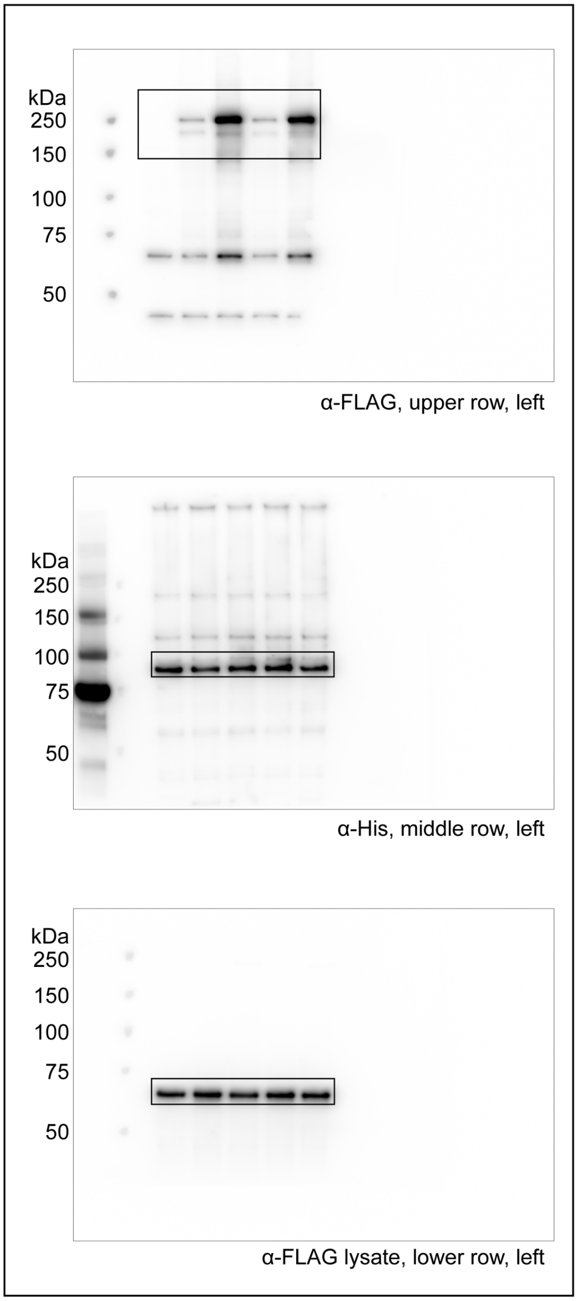


Figure 2J

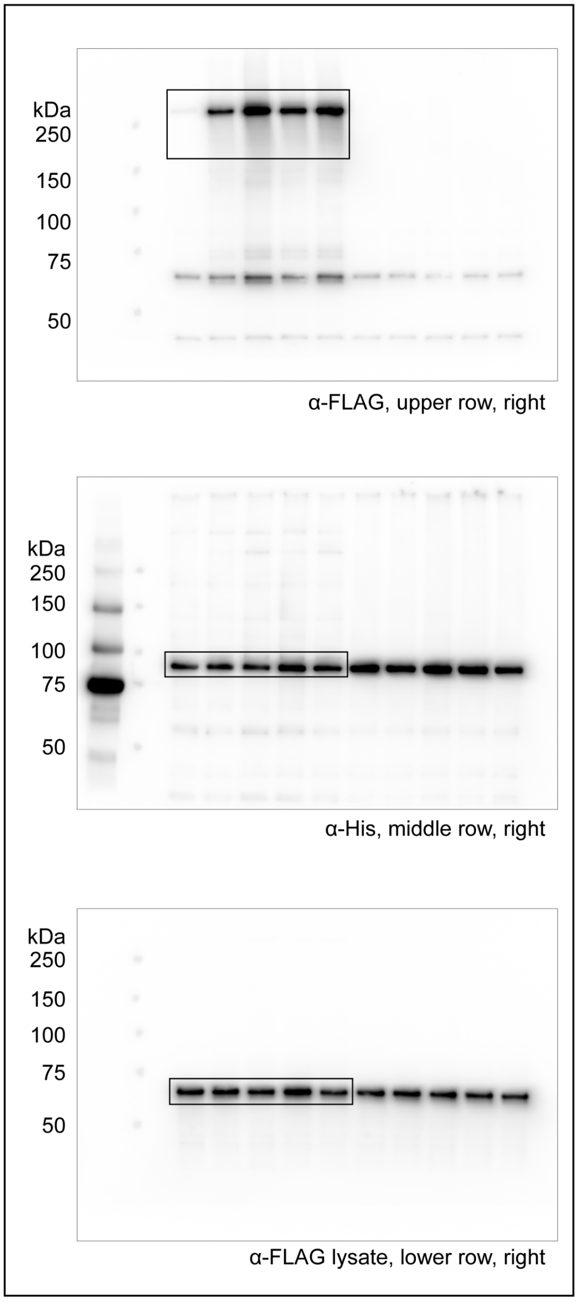

Figure 3B

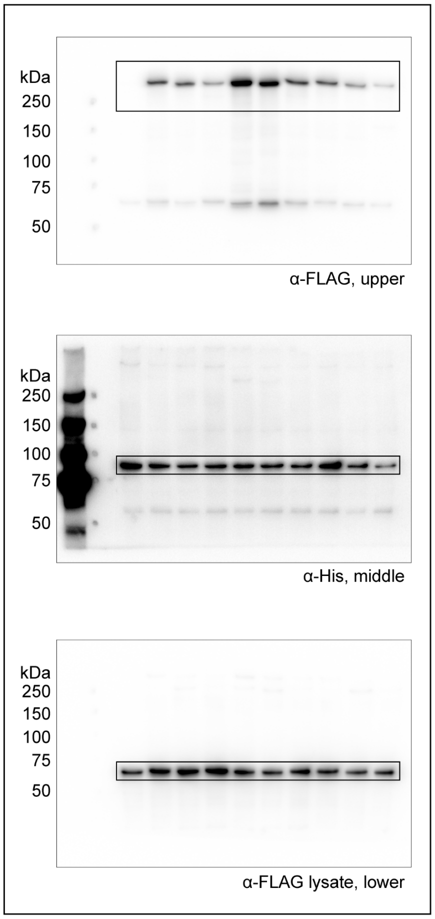


Figure 4D

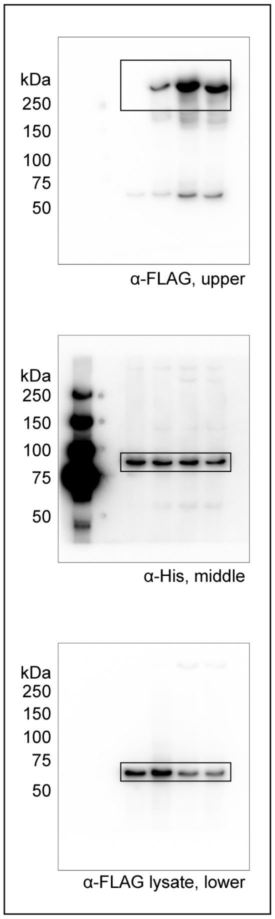


Figure 4E

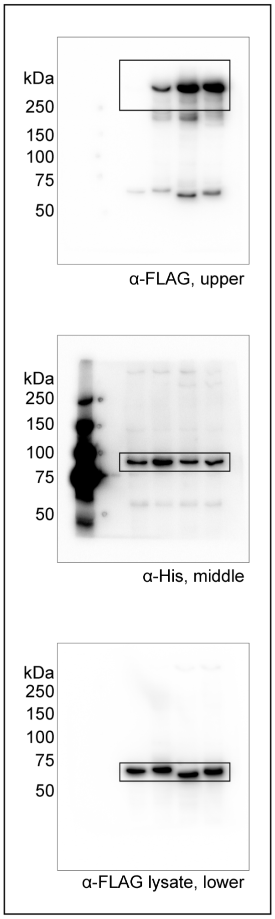


Extended Data Figure 2B

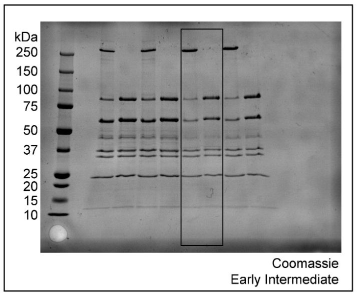


Extended Data Figure 3B

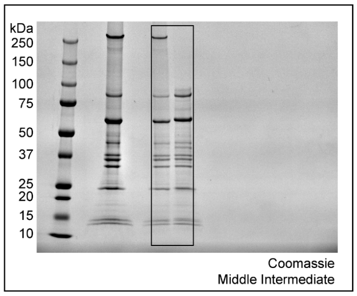


Extended Data Figure 4B

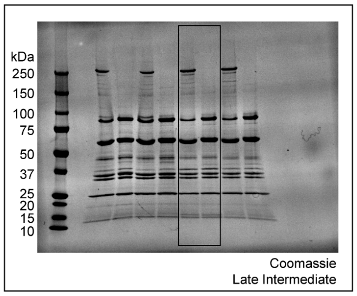


Extended Data Figure 10B

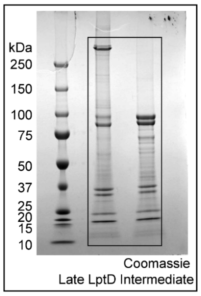

### Extended Data Figure 6E (upper)

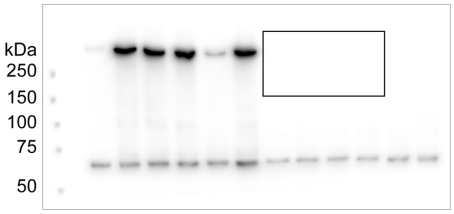


α-FLAG, upper row

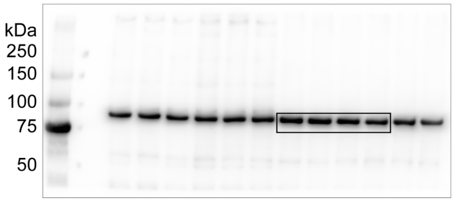


α-His, middle row

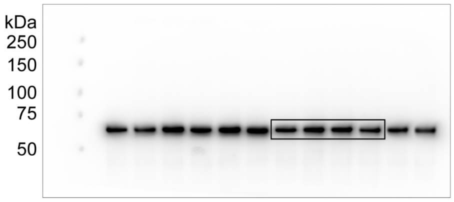


α-FLAG lysate, lower row

### Extended Data Figure 6E (lower)

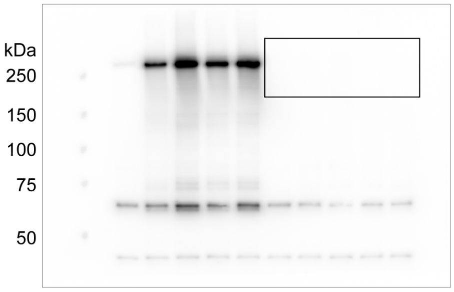


α-FLAG, upper row

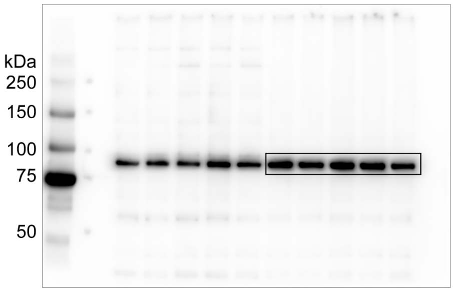


α-His, middle row

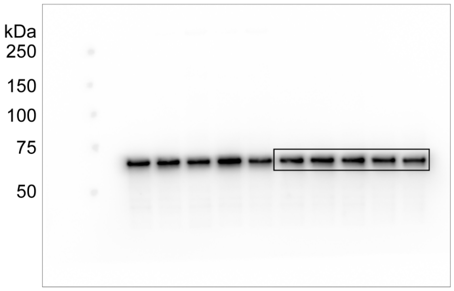


α-FLAG lysate, lower row

| Plasmid | Description | Source |
|---|---|---|
| pBDT513 | pTrc99a/8×His–BamA(D512C)–BamB–BamC–BamD–BamE | This study |
| pBDT422 | pBAD33/2×Strep–BamA(Δ172-421/Y637C) | This study |
| pBDT382 | pTrc99a/BamA(D512C)–BamB–BamC–BamD–BamE–8×His | This study |
| pBDT393 | pBAD33/2×Strep-BamA(Δ172-421/G597C) | This study |
| pBDT517 | pBAD33/2×Strep-BamA(Δ172-421/I458C) | This study |
| pBDT505 | pZS21/6×His–BamA | This study |
| pBDT374 | pZS21/6×His–BamA(D512C) | [37] |
| pBDT507 | pZS21/6×His–BamA(I806C) | This study |
| pBDT510 | pTrc99a/3×FLAG-BamA(Δ172-421/A710C) | This study |
| pBDT523 | pTrc99a/3×FLAG-BamA(Δ172-421/M711C) | This study |
| pBDT511 | pTrc99a/3×FLAG-BamA(Δ172-421/A712C) | This study |
| pBDT500 | pTrc99a/3×FLAG-BamA(Δ172-421/G636C) | This study |
| pBDT501 | pTrc99a/3×FLAG-BamA(Δ172-421/Y637C) | [37] |
| pBDT502 | pTrc99a/3×FLAG-BamA(Δ172-421/G638C) | This study |
| pBDT503 | pTrc99a/3×FLAG-BamA(Δ172-421/D639C) | This study |
| pBDT553 | pTrc99a/3×FLAG-BamA(Δ172-421/F572C) | [37] |
| pBDT378 | pTrc99a/3×FLAG-BamA(Δ172-421/G597C) | This study |
| pBDT554 | pTrc99a/3×FLAG-BamA(Δ172-421/L613C) | [37] |
| pBDT572 | pTrc99a/3×FLAG-BamA(Δ172-421/Δ601-607/F572C) | This study |
| pBDT573 | pTrc99a/3×FLAG-BamA(Δ172-421/Δ601-607/G597C) | This study |
| pBDT574 | pTrc99a/3×FLAG-BamA(Δ172-421/Δ601-607/L613C) | This study |
| pBDT586 | pTrc99a/3×FLAG-BamA(Δ172-421/Δ430-439/F572C) | This study |
| pBDT587 | pTrc99a/3×FLAG-BamA(Δ172-421/Δ430-439/G597C) | This study |
| pBDT588 | pTrc99a/3×FLAG-BamA(Δ172-421/Δ430-439/L613C) | This study |
| pBDT514 | pTrc99a/3×FLAG-BamA(Δ172-421/I458C) | [37] |
| pBDT534 | pTrc99a/3×FLAG-BamA(Δ172-421/Δ447-454/I458C) | This study |
| pBDT536 | pTrc99a/3×FLAG-BamA(Δ172-421/Δ430-439/I458C) | This study |
| pBDT538 | pTrc99a/3×FLAG-BamA(Δ172-421/Δ537-563/I458C) | This study |

| | | |
|---|---|---|
| pBDT584 | pTrc99a/3×FLAG-BamA(Δ172-421/Δ495-504/I458C) | This study |
| pTS1293 | pCDFDuet/LptE/2×Strep–LptD(ΔD330/I259C) | This study |

**Supplementary Table 1: Plasmids used in this study**

| Strain | Genotype | Source |
| --- | --- | --- |
| NovaBlue | *end*A1 *hsd*R17 ($r_K^-$, $m_K^+$) *sup*E44 *thi*-1 *rec*A1 *gyr*A96 *rel*A1 *lac* F′[*pro*A⁺ B⁺ *lac*$I^q$ ZΔM15::Tn*10*] | Novagen |
| BL21 (DE3) | *fhuA2 [lon] ompT gal (λ DE3) [dcm] ΔhsdS* | Novagen |
| MC4100 | $F^-$ *araD139 Δ(argF-lac)U169 rpsL150 relA1 flbB5301 deoC1 ptsF25 rbsR thi* | [37] |

**Supplementary Table 2: Strains used in this study**

| Description | Sequence (5' to 3') |
|---|---|
| BamA β1 Gibson F | cttcaactttggtattggttacgg |
| BamA β1 Gibson R | ccgtaaccaataccaaagttgaag |
| BamA β16 Gibson F | catcggtaaaacctggtaagtgg |
| BamA β16 Gibson R | ccacttaccaggttttaccgatg |
| BamA(G597C) F | caacctgacctgcaaagtgacc |
| BamA(G597C) R | ggtcactttgcaggtcaggttg |
| BamA(I806C) F | gttccagtttaactgcggtaaaacctg |
| BamA(I806C) R | tgttctgccttgtctccatcgtac |
| BamA(A710C) F | gcggtaactgcatggcggttg |
| BamA(A710C) R | ctacagcatcatccgatttacacag |
| BamA(M711C) F | gtaacgcctgcgcggttgcc |
| BamA(M711C) R | cgcctacagcatcatccgatttac |
| BamA(A712C) F | gtaacgccatgtgcgttgccag |
| BamA(A712C) R | cgcctacagcatcatccgatttac |
| BamA(G636C) F | ccgctggtgctatggtgatgg |
| BamA(G636C) R | gtacgccccagaacaaccc |
| BamA(G638C) F | gggttattgcgatggtttaggcgg |
| BamA(G638C) R | cagcgggtacgccccag |
| BamA(D639C) F | ggttatggttgcggtttaggcggc |
| BamA(D639C) R | ccagcgggtacgcccc |
| BamA(Δ601-607) F | tactacaaagtgacgttagacacgg |
| BamA(Δ601-607) R | ggtcactttaccggtcaggttg |
| BamA(Δ430-439) F | ttccaggctggtgtgcagc |
| BamA(Δ430-439) R | accaaagttgaagctaccggtg |
| BamA(Δ447-454) F | gctgttggtatcaacgggacc |
| BamA(Δ447-454) R | ctgctgcacaccagcctg |
| BamA(Δ537-563) F | agcttcaaaacggacgacttcacg |

| | |
|---|---|
| BamA(Δ537-563) R | cagggagttatgtacataacccag |
| BamA(Δ495-504) F | cttctataatgacttccagaccaacaagagttatgg |
| BamA(Δ495-504) R | ccataactcttgttggtctggaagtcattatagaag |

**Supplementary Table 3: Oligonucleotides used in this study**
